# Septins regulate cytokinesis and multicellular development in the closest living relatives of animals

**DOI:** 10.64898/2026.04.05.716596

**Authors:** Michael Carver, Nicole King

## Abstract

Septins are cytoskeletal proteins that regulate cytokinesis in fungi and animals, yet their functions in choanoflagellates — the closest living relatives of animals — have remained unknown. *Salpingoeca rosetta,* a choanoflagellate that switches between unicellular and multicellular forms, encodes four septins closely related to animal and fungal septins. CRISPR/Cas9-mediated disruption of *S. rosetta* septins revealed that a subset regulate cell size, with two mutants exhibiting an elevated frequency of oversized cells and one exhibiting smaller cells. Three of the four septins were required for proper rosette colony development, while two also regulated rosette structural integrity. Characterization of *Sros_septA*, which showed the strongest phenotype, revealed a role in cytokinesis: mutant cells exhibited late-stage cytokinesis failure, resulting in enlarged, multinucleated cells. Cytokinesis failure rate increased in uninduced *Sros_septA* mutant cells and was further elevated upon rosette induction, suggesting that the multicellular context places heightened demands on the septin cytoskeleton. Endogenously tagged *Sros*_SeptA dynamically redistributed from the basal pole in interphase cells to the cleavage furrow and nascent intercellular bridge during cell division. These findings identify septins as regulators of cytokinesis and multicellular development in *S. rosetta* and offer a framework for exploring how cell division regulation contributed to the emergence of animal multicellularity.

**Significance Statement:**

- Septins are cytoskeletal proteins that regulate cell division in fungi and animals, but their functions in choanoflagellates – the closest living relatives of animals – were unknown.
- Using CRISPR/Cas9 gene editing in *Salpingoeca rosetta*, we show that septins regulate both cell size and multicellular colony development. SeptA, whose gene disruption produced the strongest phenotype, localizes dynamically to the cleavage furrow and regulates cytokinesis, with cell size and division defects that are exacerbated during multicellular rosette development.
- These findings raise the possibility that elaboration of the extracellular matrix during animal origins imposed new mechanical demands on dividing cells, linking the evolution of cell adhesion to the evolution of cytokinetic regulation.

## INTRODUCTION

Septins, cytoskeletal proteins that regulate cytokinesis in fungi and animals (Hartwell *et al*., 1974; Neufeld and Rubin, 1994), have been hypothesized to contribute to multicellular development in choanoflagellates, the closest living relatives of animals (Fairclough *et al*., 2013). Although septins are found across eukaryotic diversity (Momany *et al*., 2001; Kinoshita, 2003; Shuman and Momany, 2022; Delic *et al*., 2024), they have been studied primarily in bilaterian animals and yeast. There, septins assemble into hetero-oligomeric filaments and higher-order structures that scaffold diverse factors at sites of dynamic membrane curvature (McMurray *et al*., 2011). For example, septins localize to the cleavage furrow and regulate furrow ingression, intercellular bridge formation, and abscission (Kinoshita *et al*., 2002; Surka *et al*., 2002; Estey *et al*., 2010; Founounou *et al*., 2013; Karasmanis *et al*., 2019; Hümpfer *et al*., 2025).

Due to their phylogenetic position, choanoflagellates promise to provide insights into the ancestry of animal septins (King *et al*., 2003, 2008; Fairclough *et al*., 2013; Richter *et al*., 2018). Choanoflagellates are aquatic microbial eukaryotes with a collar complex comprising an apical flagellum surrounded by microvilli (Dayel and King, 2014; Leadbeater, 2015). Under standard growth conditions, the model choanoflagellate *Salpingoeca rosetta* forms a mixture of single-celled “slow swimmers” and “chain” colonies in which cells are connected through fine intercellular bridges (Dayel *et al*., 2011). In response to environmental cues, including bacterial sulfonolipids (Alegado *et al*., 2012; Woznica *et al*., 2016) and algal polysaccharides (Perotti *et al*., 2024), *S. rosetta* instead develops into spherical “rosette” colonies in which the component cells are strongly attached through a combination of intercellular bridges, filopodia, and a shared extracellular matrix (ECM) (Dayel *et al*., 2011; Levin *et al*., 2014; Larson *et al*., 2020). Experimental control of *S. rosetta* multicellularity is one of its strengths as a model for animal origins (Booth and King, 2022).

An earlier study found that *S. rosetta* encodes four septins (renamed *Sros_septA*, *Sros_sept6*, *Sros_septB*, and *Sros_sept9* in this study; Figure 1A), and reported that their expression was upregulated in multicellular forms (Fairclough *et al*., 2013), although analyses included here differ on this point. Subsequent work using transient expression of two fluorescently tagged *S. rosetta* septins found that they localized to the basal pole of single cells and cells in rosettes (Booth *et al*., 2018). However, septin function in choanoflagellates remained unclear.

**Figure 1.**
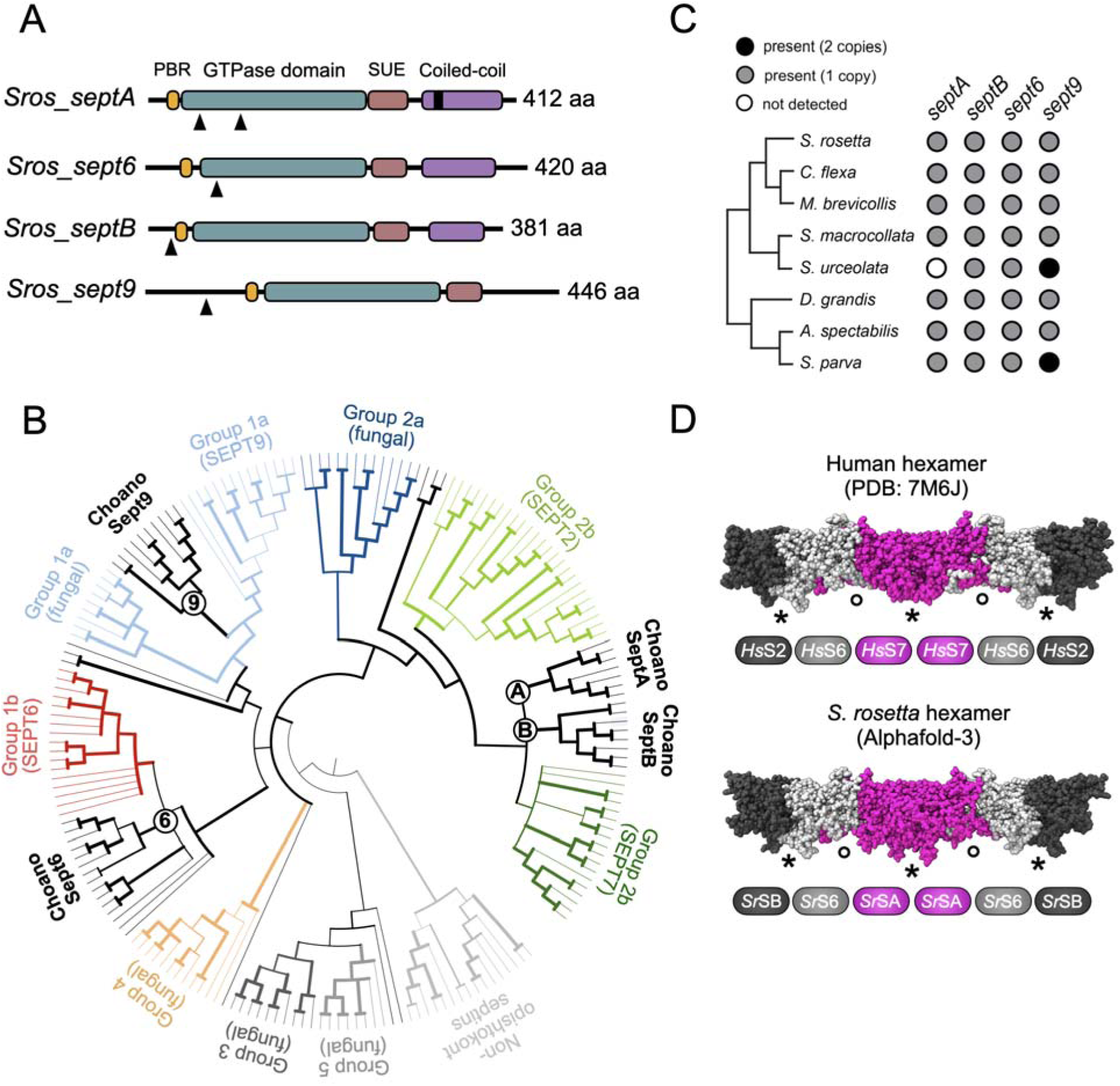
Phylogenetic relationships, conservation of protein domain architectures, and predicted hexamer assembly of *S. rosetta* septins. (A) *S. rosetta* encodes four septins – *Sros*_*septA*, *Sros*_*sept6*, *Sros*_*septB*, and *Sros*_*sept9 –* each containing a polybasic region (PBR; orange), a GTPase domain (teal), and septin-unique element (SUE; salmon). A C-terminal coiled-coil domain (purple) is present in *Sros*_*septA*, *Sros*_*sept6*, and *Sros*_*septB*, while an amphipathic helix domain (black rectangle) is present in *Sros*_*septA*. Black arrowheads indicate CRISPR-generated truncation sites (Figure S2B; Booth and King, 2020). (B) Choanoflagellate septins are closely related to animal and fungal septins. Choanoflagellate Sept9 clusters with the animal SEPT9 clade (within Group 1a; light blue), while choanoflagellate Sept6 forms a polytomy with animal SEPT6 proteins (red), septins from other close animal relatives, and the *C. elegans* septin *UNC-61* (Group 1b) (Shuman and Momany, 2022). Choanoflagellate SeptA and SeptB form a polytomy with animal SEPT7 within the broader SEPT2/7 animal clade (Group 2b; dark green). A maximum-likelihood phylogeny is shown of predicted septin protein sequences from diverse opisthokonts and selected non-opisthokont taxa (File S1). Branch width scales with UFboot support and all nodes with less than 80% bootstrap support were collapsed. Branch lengths do not scale with evolutionary distance in this rendering. See Figure S1A for full annotated version of this phylogeny. (C) Choanoflagellate septins analyzed in this study belong to four families – *septA*, *septB*, *sept6*, and *sept9* – each broadly distributed across choanoflagellate diversity. The presence of two genes (black circle), one gene (grey circle), or no detected genes (white circle) is indicated. Choanoflagellate septin families were defined based on the septin phylogeny in this study (Figure 1B and Figure S1A) and are shown within a composite maximum likelihood phylogeny of choanoflagellates, derived from Carr *et al*. (2017) with the addition of *C. flexa* from Brunet *et al*. (2019). (D) *S. rosetta* septins are predicted to form hexamers resembling those of humans (shown) and other opisthokonts. Human septins within the cryo-EM structure of the human septin hexamer (Mendonça *et al*., 2021): SEPT2 (*Hs*S2; dark grey), SEPT6 (*Hs*S6; light grey), and SEPT7 (*Hs*S7; magenta). *S. rosetta* septins modeled by AlphaFold-3: *Sros_*SeptB (*Sr*SB; dark grey), *Sros_*Sept6 (*Sr*S6; light grey), and *Sros_*SeptA (*Sr*SA; magenta), with termini trimmed to match the residue coverage of the solved human structure (Materials and Methods). Asterisks denote N- to C-terminal interfaces; circles denote GTP-binding domain interfaces. Confidence scores are given in Figure S1B.

Here, we set out to understand septin function in *S. rosetta*. We find that *Sros*_*septA*, *Sros_sept6*, and *Sros_*sept9 regulate cell size in opposing directions (*Sros*_*septA* and *Sros_sept6* losses increase cell size; *Sros_sept9* loss decreases it) and all three are required for proper rosette assembly, while *Sros*_*septA* and *Sros*_*sept6* additionally regulate rosette integrity. Further characterization of *Sros_septA* revealed a role in cytokinesis, with associated cell size and division defects that become more pronounced during rosette development. Consistent with its role as a regulator of cytokinesis, *Sros*_SeptA localization is dynamic throughout the cell cycle, localizing to the basal pole in interphase cells and to the cleavage furrow and nascent intercellular bridge during cytokinesis. Together, these findings suggest a role for septins at the interface of cytokinesis regulation and the evolution of animal multicellularity.

## RESULTS

### *S. rosetta* septins are closely related to animal and fungal septins and are predicted to assemble into hexamers

Choanoflagellate genomes and transcriptomes published after the 2013 study of *S. rosetta* septins (del Campo and Ruiz-Trillo, 2013; Richter *et al*., 2018; Brunet *et al*., 2019; López-Escardó *et al*., 2019; Hake *et al*., 2024) prompted us to construct an updated phylogeny of septins using protein sequences from diverse choanoflagellates and other eukaryotes, including early-branching animals and fungi. Consistent with Fairclough *et al*. (2013), *Sros*_Sept9 was recovered as sister to animal SEPT9 sequences (Figure 1B and S1A), and *Sros_*Sept6 fell within an unresolved polytomy that included the animal SEPT6 clade, septins from other close animal relatives, and the *Caenorhabditis elegans* septin *UNC-61*. The two remaining *S. rosetta* septins – previously called Septin2 and Septin7 – nested within the broader animal SEPT2/SEPT7 radiation (Kinoshita, 2003) but could not be resolved relative to one another or the animal SEPT7 group, forming a three-way polytomy. To avoid implying specific orthology that is not supported by our current phylogenetic analysis, we renamed *S. rosetta* Septin2 to *Sros_*SeptA and Septin7 to *Sros*_SeptB. Furthermore, each of the four *S. rosetta* septins cluster within clades that include septins from across choanoflagellate diversity, indicating broad conservation of these septin families (Figure 1C).

To investigate whether *S. rosetta* septins can assemble into hexameric structures similar to those found in animals and fungi (Sirajuddin *et al*., 2007; Bertin *et al*., 2008), we predicted their interactions using AlphaFold-3 (Abramson *et al*., 2024). *Sros_*SeptA, *Sros_*Sept6, and *Sros_*SeptB were predicted to form a hexamer similar to the reported hexamers from humans (Figure 1D and S1B; Mendonça *et al*., 2021) and other opisthokonts. The AlphaFold-3 analysis placed *Sros_*SeptA in the position typically occupied by animal SEPT7, while *Sros_*SeptB was placed in the position typically occupied by animal SEPT2. These data are consistent with conservation in *S. rosetta* of the hexameric septin assembly found in animals and yeast. Currently, there is no published structure of the SEPT9-containing septin octamer, although the subunit arrangement has been inferred experimentally (Kim *et al*., 2011). Furthermore, inclusion of SEPT9 in AlphaFold-3 predictions of human septin interactions did not recapitulate this inferred octameric architecture. Therefore, comparisons of hexameric structures were used for our analyses.

### Multiple septins regulate cell size and rosette development

The first analyses of *S. rosetta* septins suggested that septin gene transcription was elevated in colonial cells (rosettes and chains; Figure 2A) relative to solitary cells (Fairclough *et al*., 2013). Because of subsequent improvements to cell state regulation protocols for *S. rosetta*, we reanalyzed septin transcription using a recently published RNA-seq dataset (Leon *et al*., 2025). In this dataset, four populations of cells were compared: (1) slow swimmers and chains, which naturally co-occur under standard growth conditions, (2) rosettes, (3) cultures enriched for single-celled fast swimmers, and (4) thecates, a single-celled sedentary cell state. Transcription of all four septins was reduced in thecate cells relative to slow swimmers and chains, and *Sros*_*septB* was additionally downregulated in fast swimmers (Figure S2A). However, transcription of *Sros*_*septA*, *Sros*_*sept6*, and *Sros*_*sept9* in slow swimmers and chains was comparable to that in both rosette cultures and fast swimmer cultures. Together, these data indicate that overall septin expression is not specifically upregulated in colonial cell types.

**Figure 2.**
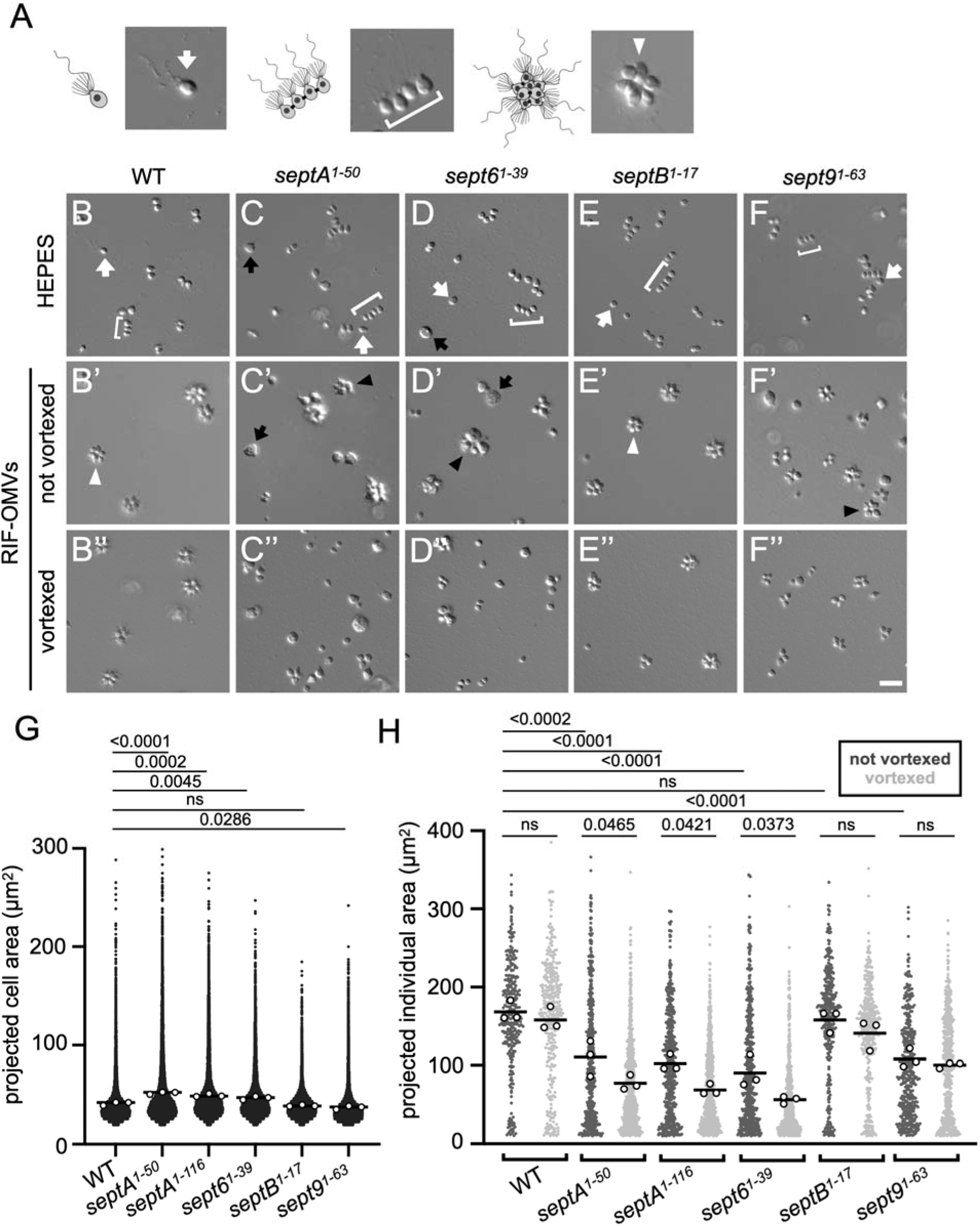
Septins regulate cell size and the structural integrity of rosettes. (A) The life history of *S. rosetta* includes single cells (left), chains (middle), and rosettes (right). (B – F’’) Defects in cell size and rosette morphology in septin mutants. (B – F) Wild-type (B) and septin mutants (C – F) grown with carrier control (HEPES–KOH, pH 7.5) produced slow swimmers (white arrows) and chains (brackets). *Sros*_*septA^1-50^* (C) and *Sros*_*sept6^1-39^* cultures (D) frequently contained larger cells (black arrows). (B’ – F’) After treatment with RIF-OMVs, wild-type (B’) and *Sros*_*septB^1-17^* cultures (E’) produced compact rosettes of comparably-sized cells (white arrowheads), whereas *Sros*_*septA^1-50^*, *Sros*_*sept6^1-39^*, and *Sros*_*sept9^1-63^* strains (C’, D’, and F’) produced abnormal rosettes with both normal and comparatively larger cells (black arrowheads). (B’’ – F’’) Wild-type (B’’), *Sros_septB^1-17^* (E’’), and *Sros_sept9^1-^*^63^ rosettes (F’’) maintained their structural integrity after vortexing, while *Sros_septA^1-50^*(C’’) and *Sros_sept6^1-39^* cultures (D’’) contained colonies with fewer cells and more single cells after vortexing, indicating reduced structural integrity. Scale bar, 20 μm. (G) When grown without RIF-OMVs, *Sros*_*septA^1-50^* and *Sros*_*sept6^1-39^* strains exhibited significantly increased mean cell sizes compared with wild-type, while the mean cell size in *Sros_sept9^1-63^*was reduced. Black dots represent projected cell area of individual cells pooled from three biological replicates (18,382 – 23,313 total cells); white dots represent means per replicate; bars denote means across replicates. Statistics performed on replicate means (n = 3): one-way ANOVA with Dunnett’s multiple comparisons test. Further analysis of size distributions is provided in Figure S2F. (H) Septin mutants showed defects in both rosette assembly and rosette integrity. After RIF-OMV treatment, *Sros_septA^1-50^*, *Sros_septA^1-116^, Sros_sept6^1-39^,* and *Sros_sept9^1-63^* formed smaller structures compared with wild-type, indicating defects in rosette assembly (“rosette assembly defect”). Upon vortexing, the size of individuals in *Sros*_*septA^1-50^*, *Sros_septA^1-116^,* and *Sros*_*sept6^1-39^* significantly decreased (“rosette integrity defect”). Dark grey and light grey dots represent projected area of individual cells or multicellular morphs pooled from three biological replicates for unvortexed and vortexed conditions, respectively (318 – 723 total individuals); white dots represent means per replicate; bars denote means across replicates. Statistics performed on replicate means (n = 3): two-way ANOVA with Dunnett’s multiple comparisons test for between-strain and Šidák’s test for within-strain comparisons.

To investigate septin function in *S. rosetta*, we introduced a premature stop codon near the 5’ end of each septin gene (Figures 1A and S2B) using CRISPR/Cas9-mediated genome editing. The resulting mutants – *Sros_septA^1-50^, Sros_septA^1-116^, Sros_septB^1-17^, Sros_sept6^1-39^,* and *Sros_sept9^1-63^* – were named based on the number of N-terminal amino acids remaining after each truncation. Because *Sros_septA^1-50^*consistently exhibited more severe phenotypes than *Sros_septA^1-116^*across all assays, *Sros_septA^1-50^* is shown as the representative *septA* allele in Figures 2–4; *Sros_septA^1-116^* images and quantification are provided in Figure S2D and included in most summary graphs. *Sros_septA^1-50^* and *Sros_sept6^1-39^* each exhibited a slight proliferation defect during log phase (Figure S2C), while other septin mutants (including the other *Sros*_*septA* allele, *Sros_septA^1-116^*) matched the proliferation dynamics of wild-type cells.

**Figure 3.**
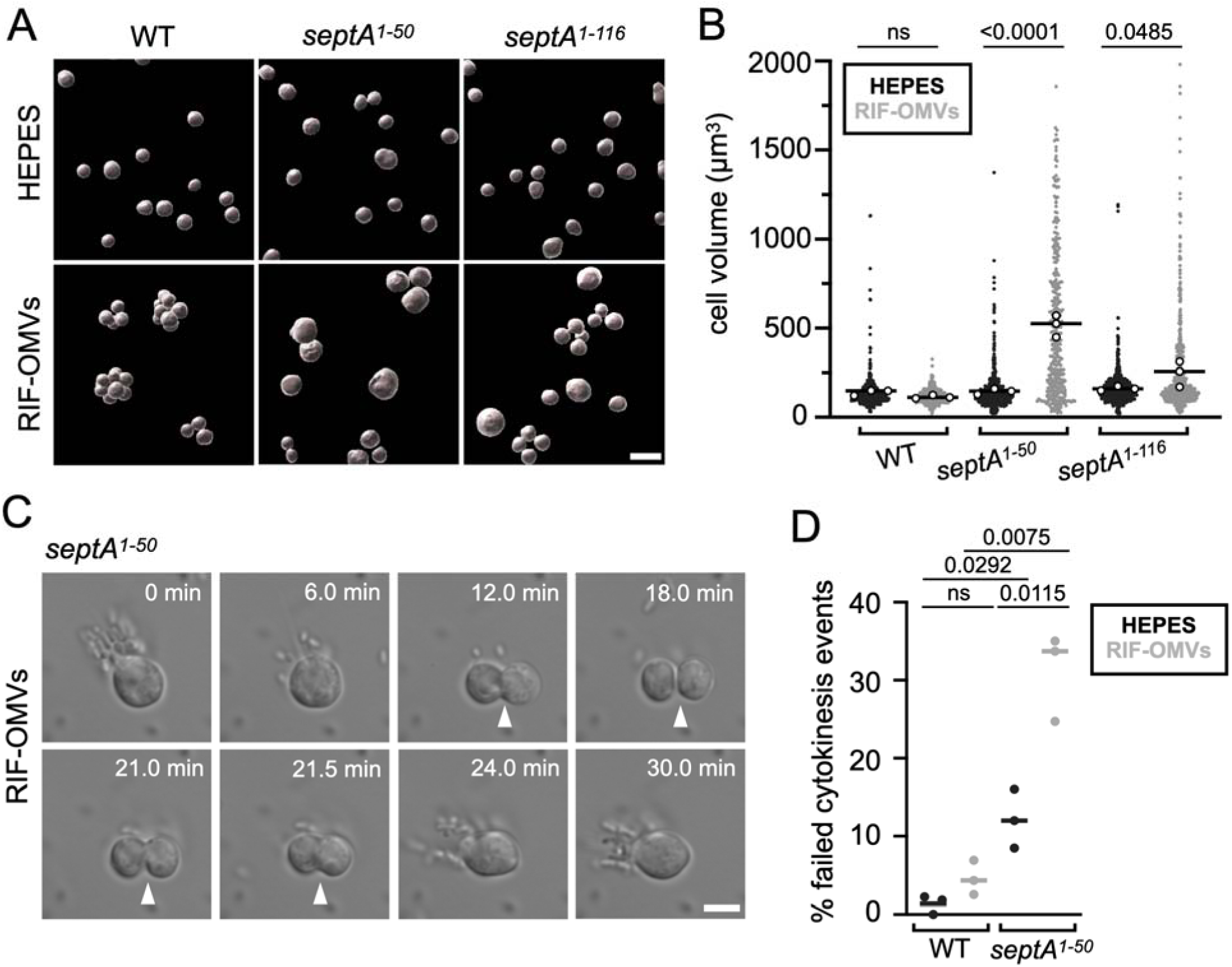
Rosette induction in *Sros*_*septA* mutants leads to increased cell size and cytokinesis defects. (A) Cell segmentation enables quantification of cell volume in single and rosette cells (Figure S3B). Scale bar, 10 μm. (B) Mean cell volume increased significantly after rosette induction in *Sros*_*septA* mutants, but not in wild-type. Black and grey dots represent cell volumes of individual cells pooled from three biological replicates (359 – 600 cells total) for HEPES and RIF-OMV treatments, respectively; white dots represent means per replicate; bars denote means across replicates. Statistics performed on replicate means (n = 3): two-way ANOVA with Šidák’s multiple comparisons test. (C) Time-lapse imaging of a representative *Sros*_*septA^1-50^* cell after rosette induction reveals late-stage cytokinesis failure (Video S2). White arrowheads indicate cleavage furrow ingression and cell fusion during cytokinesis progression and collapse (12.0–21.5 min). Scale bar, 5 μm. (D) RIF-OMV treatment increases the rate of cytokinesis failure in *Sros*_*septA^1-50^* cells. In wild-type cells, cytokinesis failure was rare and did not differ significantly between untreated and RIF-OMV-treated conditions. In contrast, *Sros*_*septA^1-50^* exhibited an elevated baseline failure rate relative to wild-type, which increased more than twofold upon RIF-OMV treatment. Dots represent means per replicate; bars denote means across three biological replicates (166-208 cell division events analyzed in total). Statistics performed on replicate means (n = 3): unpaired t-tests with Welch’s correction for each pairwise comparison.

**Figure 4.**
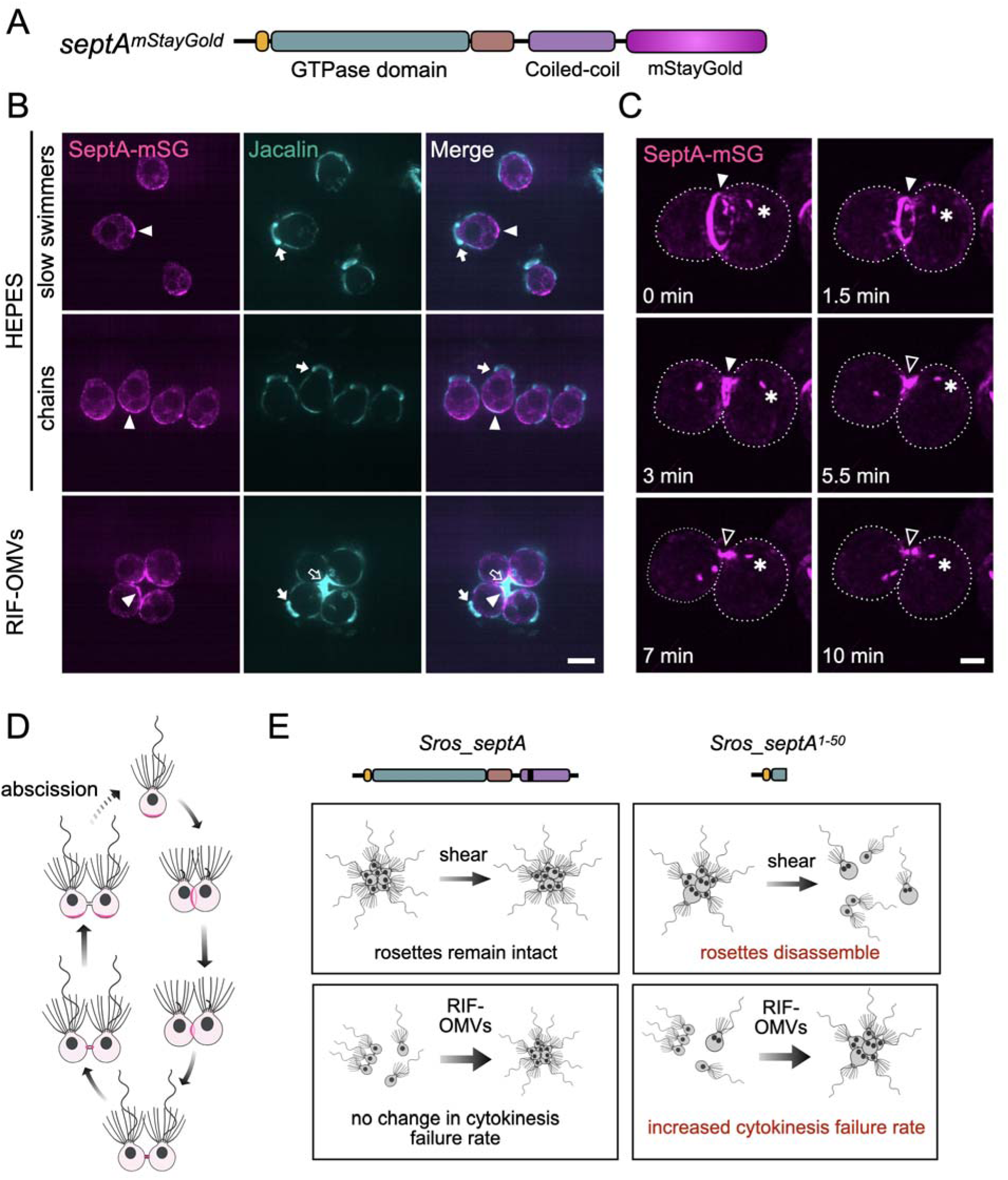
SeptA localizes to the basal pole in interphase *S. rosetta* cells and to the cleavage furrow and nascent intercellular bridge in dividing cells. (A) Domain structure of endogenously tagged SeptA-mStayGold (SeptA-mSG). (B) SeptA-mSG is distributed throughout the cytoplasm and enriched at the basal pole in interphase single cells (top row), cells in chains (middle row), and cells in rosettes (bottom row). The lectin Jacalin (teal) marks the collar base (filled arrow) and the ECM of rosettes (open arrow). HEPES served as carrier control for untreated cultures (yielding slow swimmers and chains); RIF-OMVs induced rosettes. Arrowheads mark basal poles of cells. Scale bar, 5 μm. (C) SeptA-mSG accumulates in a ring at the cleavage furrow (filled arrowhead). As cleavage progresses, the ring constricts and SeptA-mSG becomes enriched at the nascent intercellular bridge (open arrowhead), resolving into two puncta after several minutes. Dotted lines outline cell boundaries. Scale bar, 1 μm. Dynamic SeptA-mSG puncta of unknown function were also detected (asterisk). (D) Proposed model for SeptA localization during the cell cycle. In interphase, SeptA (magenta) localizes throughout the cell body and is enriched at the basal pole. During cytokinesis, the flagellum retracts and SeptA redistributes to the cleavage furrow, then to the nascent intercellular bridge. After abscission, SeptA returns to the basal pole. (E) Summary of phenotypes associated with *Sros_septA^1-50^.* Wild-type cells show no change in cytokinesis failure rate following RIF-OMV treatment and form shear-resistant rosettes. In contrast, *Sros_septA^1-50^* mutants exhibit increased cytokinesis failure after rosette induction, resulting in enlarged cells, and form rosettes that disassemble upon shear.

We next checked whether septin loss altered the morphology of *S. rosetta* slow swimmers and chains (Figure 2A). Cultures of all five septin mutants contained mixtures of single cells and chains that resembled those of wild-type cultures (Figure 2B-F, Figure S2D). However, *Sros_septA^1-50^*, *Sros_septA^1-116^*, and *Sros_sept6^1-39^* had a small increase in the frequency of oversized cells relative to wild-type (Figure 2C; Figure S2D and S2F), which increased mean cell size by 1.25-fold, 1.19-fold, and 1.13-fold in *Sros_septA^1-50^*, *Sros_septA^1-116^*, and *Sros_sept6^1-39^*, respectively (Figure 2G and Table S2). In contrast, mean cell size decreased by 0.90-fold in *Sros_sept9^1-63^* and was not significantly different in *Sros_septB^1-17^*.

Rosette development in *S. rosetta* can be induced by treatment with sulfonolipids called Rosette Inducing Factors that are released in outer membrane vesicles (RIF-OMVs) from the bacterium *Algoriphagus machipongonensis* (Fairclough *et al*., 2010; Woznica *et al*., 2016). After treatment with RIF-OMVs, *Sros_septA^1-50^*, *Sros_septA^1-116^, Sros_sept6^1-39^*, and *Sros_sept9^1-63^* cultures exhibited morphological defects (Figure 2C’, D’, and F’, and Figure S2D), with some rosettes having heterogeneous cell sizes and disrupted organization relative to wild-type, as well as an increased number of non-rosette single cells. As a proxy measure of rosette assembly, we quantified the mean areas of individuals (single cells, doublets, triplets, or rosettes) after RIF-OMV induction and found that the mean individual area was significantly reduced across all morphological classes in *Sros_septA^1-50^*, *Sros_septA^1-116^, Sros_sept6^1-39^*, and *Sros_sept9^1-63^*cultures, indicating a reduction in the sizes of rosettes (Figure 2H and S2G; Table S3). In contrast, RIF-OMV-treated *Sros*_*septB^1-17^* showed no significant size reduction and formed rosettes that resembled wild-type (Figure 2H and 2E’; Table S3).

Wild-type rosettes exhibit robust structural integrity and resistance to shear (Levin *et al*., 2014). To assess whether septins affect rosette integrity, we vortexed wild-type and mutant cultures after rosette induction and quantified the projected area of the resulting individuals (Wetzel *et al*., 2018; Xu *et al*., 2022). In wild-type cultures, cells in rosettes remained attached; vortexing caused no significant reduction in mean projected individual area (Figure 2B’’ and H; Table S4). Similarly, the mean projected area of individuals in *Sros_septB^1-17^*and *Sros_sept9^1-63^* cultures induced with RIF-OMVs did not significantly change after shear (Figure 2E’’, 2F’’ and 2H; Table S4). In contrast, *Sros*_*septA^1-50^, Sros*_*septA^1-116^,* and *Sros*_*sept6^1-39^* cultures experienced significant disassembly (Figure 2C’’ and D’’; Figure S2G). Because ECM production is important for proper rosette development, we hypothesized that septin mutants with compromised rosette integrity might exhibit ECM defects. However, staining of *Sros_septA^1-50^*rosettes with two markers of ECM, the lectin Jacalin (Figure S3A) and an antibody to the ECM protein Rosetteless (Figure S3A’) (Levin *et al*., 2014; Wetzel *et al*., 2018), revealed that ECM levels were qualitatively similar to those in wild-type rosettes. This suggests that the loss of rosette integrity does not stem from an obvious loss of ECM secretion, although these data do not rule out subtler defects in ECM composition and organization.

### Rosette induction in *Sros_septA* mutants results in increased cell size and cytokinesis defects

To better understand the connection between septin function and rosette development, we focused the remainder of our analyses on the *Sros_septA* mutants, which exhibited the strongest defects in cell size regulation and rosette development. We first asked whether cell size differed in *Sros_septA* mutants before and after rosette induction. Because the projected area of each cell cannot be easily measured in rosettes, we instead used cell segmentation to quantify cell volume (Figure 3A and S3B). In wild-type cultures, untreated (i.e. slow swimmer and chain) and RIF-OMV-treated (i.e. rosette) conditions did not significantly differ in mean cell volume (Figure 3B; Table S5). In contrast, RIF-OMV treatment of *Sros_septA^1-50^* led to a 3.5-fold increase in mean cell volume over untreated *Sros_septA^1-50^* cells, while *Sros_septA^1-116^* cell volume increased by more than 1.5-fold (Table S5). Whether this volume increase is selective for cells incorporated into rosettes, or also occurs in the isolated single cells present in RIF-OMV-treated cultures, was not directly assessed in this experiment. This increase indicates a heightened importance of *Sros_septA* for regulating cell volume during rosette development.

Increased cell size in *S. cerevisiae* and *D. melanogaster* septin mutants is often a consequence of cytokinesis defects (Hartwell *et al*., 1973; Neufeld and Rubin, 1994). Therefore, we next examined cytokinesis in *Sros_septA^1-50^*mutants. In wild-type *S. rosetta* cells, mitosis occurs along the apical–basal axis, with daughter cells either completing abscission or remaining attached by a narrow intercellular bridge, although the relative frequency of these outcomes has not been characterized (Figure S3C and S3D, Video S1; (Dayel *et al*., 2011; Fairclough *et al*., 2013)). In *Sros_septA^1-50^*, cells that had undergone cleavage furrow ingression and appeared to have separated into daughter cells frequently fused back together, indicating a defect in a late stage of cytokinesis (Figure 3C, Video S2). In all observed failure events, furrow ingression proceeded to this apparent separation step, consistent with a specific requirement for *Sros*_SeptA during late cytokinesis rather than during earlier stages of furrow ingression. We also noted that in some *Sros_septA*¹ divisions, the cleavage plane appeared misaligned with the apical–basal cell axis, raising the possibility of a spindle positioning defect; whether this contributes to cytokinesis failure will require further investigation (Videos S2, S3). To investigate the fate of the nucleus during failed cytokinesis events in *Sros_septA^1-50^*, we generated *Sros_septA^1-50^* cells that expressed histone H2B fused to mCherry (*Sros_septA^1-^*^50^; *h2b-mCherry*) as a marker of nuclei. In these cells, the nuclei divided normally as the cell attempted cytokinesis and remained distinct following cytokinesis failure, resulting in large, multinucleated cells (Figure S3E, Video S3).

Consistent with the *Sros*_*septA^1-50^*cell volume phenotype, the rate of cytokinesis failure in *Sros*_*septA^1-50^* was elevated in uninduced conditions and increased further – more than doubling – upon rosette induction during the first ten hours following treatment. In contrast, cytokinesis failure rates in wild-type cells were low and not significantly affected by rosette induction (Figure 3D; Table S6). Together, these data suggest that *Sros*_*septA* is more important for completion of cytokinesis in rosette cells than in uninduced cells.

### Dynamic localization of *Sros*_SeptA during cytokinesis

To visualize *Sros*_SeptA *in vivo*, we adapted a recently published CRISPR-based gene editing strategy (Combredet *et al*., 2025) to allow endogenous tagging of *S. rosetta* proteins with fluorescent proteins (Materials and Methods). Using this approach, we tagged the C-terminus of *Sros*_SeptA with mStayGold, a photobleach-resistant monomeric fluorescent protein (Ivorra-Molla *et al*., 2024; Figure 4A). The monomeric form was used to minimize potential aggregation or multimerization artifacts. In addition, we tagged *Sros*_SeptA with an ALFA tag, a short epitope that can be detected using a fluorescently labeled nanobody (Götzke *et al*., 2019). Both *Sros*_SeptA-mStayGold (SeptA-mSG) and *Sros*_SeptA-ALFA were detected primarily at the basal poles of non-dividing single cells, cells in chains, and cells in rosettes, with diffuse signal observed throughout the remainder of the cytoplasm (Figure 4B; Figure S4A and B). These observations were consistent with prior observations of transiently-expressed *Sros*_SeptA-mWasabi (Booth *et al*., 2018).

To assess whether the mStayGold tag affected *Sros*_SeptA function, we examined cell size and rosette integrity in *Sros*_*septA-mSG* and *Sros*_*septA-ALFA* cells. Neither strain exhibited the cell size defect observed in *Sros_septA^1-50^*cells following induction with RIF-OMVs (Figure S4D and S4E). However, rosette integrity in *Sros*_*septA-mSG* and *Sros*_*septA-ALFA* cells was compromised relative to wild-type cells (Figure S4F), indicating that the rosette development phenotype is separable from the cell size defect in tagged *Sros*_*septA* strains.

Because *Sros*_*septA-mSG* cells did not exhibit a mutant cell size phenotype, we used this strain to analyze *Sros*_SeptA dynamics during cell division. *Sros*_SeptA-mSG in dividing cells accumulated in a ring at the cleavage furrow during furrow ingression and, in contrast to interphase cells, did not appear to be enriched at the basal pole (Figure 4C; Figure S4H; Video S4 and S5). As cytokinesis progressed, the *Sros_*SeptA-mSG ring constricted until it was largely restricted to the nascent intercellular bridge connecting the daughter cells, a pattern also detected by immunofluorescence in *Sros*_*septA-ALFA* cells (Figure S4C). We observed no clear differences in septin localization or septin ring constriction when comparing single cells and rosettes. Together, these observations suggest that *Sros*_SeptA localization is dynamic across the cell cycle in single cells and rosettes (Figure 4D).

## DISCUSSION

Despite extensive research on septin function in fungi and bilaterian animals (Spiliotis and McMurray, 2020), septin function in non-bilaterians and close relatives of animals has been largely unexplored (Fairclough *et al*., 2013; Booth *et al*., 2018). Additionally, the role that cell division regulators might have played in the emergence of multicellularity in animals remains an open question (Arakaki *et al*., 2017; Brunet and King, 2017; Chaigne and Brunet, 2022). By examining septin function in *S. rosetta*, a choanoflagellate in which multicellular development can be experimentally controlled, this study helps address both gaps. Here, we identify a role for the septin cytoskeleton in regulating cell size and cytokinesis and show that this requirement is heightened during rosette development (Figure 4E). Specifically, disruption of *Sros_septA* in rosette-induced cells results in increased cell size and an elevated rate of cytokinesis failure relative to uninduced cells, suggesting that the multicellular context influences septin function in *S. rosetta*.

One possible explanation is that ECM attachment — beginning at the first division and persisting throughout rosette morphogenesis (Larson *et al*., 2020) — introduces mechanical constraints that place additional demands on the septin cytoskeleton during cytokinesis, for example by tethering dividing cells to a shared ECM that resists the membrane remodeling required for cell division. This raises the possibility that ECM elaboration, by physically constraining dividing cells within a shared matrix, imposed new demands on cytokinetic completion, linking the evolution of cell adhesion to the evolution of cell division regulation. Alternatively, the phenotype may reflect biochemical responses to the inducing signal itself, rather than the mechanical constraints imposed by rosette architecture. A third possibility is that septins are differentially regulated in rosettes compared with uninduced cells. However, although *Sros*_SeptA dynamically redistributes throughout the cell cycle, we do not find any obvious differences in *Sros*_SeptA localization or transcription levels between single cells and rosettes.

We also find that *Sros*_*septA*, *Sros*_*sept6*, and *Sros*_*sept9* regulate rosette assembly in *S. rosetta* while *Sros*_*septA* and *Sros*_*sept6* also regulate rosette integrity, raising the possibility that septins directly regulate multicellular development (Figure 4E). While changes in cell size may secondarily affect rosette assembly or integrity, our finding that endogenously tagging *Sros*_*septA* disrupts rosette development without altering cell size suggests that septins may also act through cell size-independent mechanisms. For example, septins may help to scaffold factors at the basal pole to promote cell-ECM interaction or may influence the positioning or stability of intercellular bridges that physically connect cells within rosettes. The absence of a detectable phenotype in *Sros_septB^1-17^*, and the dispensability of *Sros_sept9* for rosette integrity, may reflect incomplete loss-of-function alleles, functional redundancy, or roles for these septins that are distinct from those of SeptA and Sept6 — questions that future work can address. Together, these findings show that the septin cytoskeleton regulates cytokinesis and multicellular development in *S. rosetta*. More broadly, our results suggest that septins represent a plausible link between cytokinesis and multicellular organization and offer a framework for exploring how cell division regulation intersected with the evolution of animal multicellularity.

## MATERIALS AND METHODS

### BLAST searches for septin genes

To re-investigate the presence of septin genes across opisthokonts, we used human septin proteins representing the four major animal septin groups as BLASTP queries: SEPT2 (NP_004395.1), SEPT6 (NP_006138.1), SEPT7 (NP_001779.2), and SEPT9 (NP_006631.1). We searched for putative septins in 18 animals (eight bilaterians, four cnidarians, one placozoan, two ctenophores, and three poriferans), eight choanoflagellates, five additional holozoan relatives (filastereans, corallochytreans, and ichthyosporeans), two early-branching holomycotans, and seven fungi (File S1). Additionally, we included septins recently reported from nine non-opisthokont species (Delic *et al*., 2024). Species were selected to (1) sample diverse choanoflagellates (Richter *et al*., 2018), (2) increase representation of close animal relatives and early-diverging animals and fungi, and (3) include established model organisms (e.g. *D. melanogaster*, *S. cerevisiae*).

Searches were performed against the EukProt database (Richter *et al*., 2022), a curated collection of predicted eukaryotic proteomes derived from genome and transcriptome assemblies, using BLASTP v2.13.0 with an E-value cutoff of 1 × 10. For *Hydra vulgaris*, *Acropora digitifera*, and *Magnaporthe oryzae*, which are not hosted on EukProt, searches were performed using the NCBI nonredundant protein database (Sayers *et al*., 2023). Candidate septins were defined as proteins containing the conserved septin GTP-binding domain (Pfam accession PF00735), identified using InterProScan (Jones *et al*., 2014), which detects Pfam domains through Hidden Markov Model-based searches performed with HMMER. For bilaterian taxa, where septin family membership has been relatively well described (Kinoshita, 2003; Shuman and Momany, 2022), only the top septin hit from each of the four BLAST searches using human septin queries was included in the phylogeny for simplicity, with the exception of *D. melanogaster* (File S1).

For non-bilaterian animals and other taxa, all recovered septin-domain proteins were included. Additionally, the top putative septin BLAST hit for all non-bilaterians was used as a query in a second BLAST search against the same proteome to identify additional paralogs. Iterative searches continued until no additional septin-domain proteins were recovered. Using this approach, a set of 201 putative septin sequences was recovered (File S1).

To further refine the dataset, sequences were filtered based on domain confidence, protein length, and domain architecture (File S1). All detected septin domain matches met the E-value cutoff of 1 × 10^-5^ (InterProScan), so we next turned to protein length. Septin proteins are typically ∼30 – 65 kDa (Mostowy and Cossart, 2012), or approximately 300 – 600 amino acids, although N-terminal extensions exist in the SEPT9-family (Kim *et al*., 2011). We therefore excluded candidate protein sequences below 200 amino acids and above 1000 amino acids, removing eight sequences. The remaining sequences were manually inspected to confirm the absence of domains inconsistent with canonical septins (e.g., DNA-binding or kinase domains). One fungal sequence containing a zinc finger DNA-binding domain was removed. The final dataset comprised 192 septin sequences (File S1) and was used for our phylogenetic analysis (Figure S1A).

### Phylogenetic trees

To infer maximum-likelihood phylogenies of septin genes (Figure 1C; Figure S1A), we aligned protein sequences using MAFFT v7 with the L-INS-i algorithm set to default parameters, trimmed with ClipKIT v1.3.0 (Steenwyk *et al*., 2025) using the *smart-gap* mode, and analyzed using IQ-TREE v3.0.1 (Wong *et al*., 2025). Ten independent maximum-likelihood searches were performed using IQ-TREE under the best-fitting substitution model (Q.PFAM+R8), each initiated from different random starting seeds. Branch support was estimated using ultrafast bootstrap approximation (UFBoot) with 1,000 replicates, and the tree with the highest log-likelihood score was selected for downstream analysis. Branches with bootstrap support below 80% were then collapsed prior to visualization. Trees were visualized using the Interactive Tree of Life (iTOL) web server (Letunic and Bork, 2021) and rooted using non-opisthokont septins as an outgroup (Delic *et al*., 2024). Sequence labels in Figure S1A correspond to the species name followed by the EukProt protein identifier, the RefSeq accession, or the identifiers for non-opisthokont septins used in Delic *et al*. (2024). The full amino acid sequences used in the phylogenetic analysis are provided in File S1. Sequence datasets, multiple sequence alignments, phylogenetic trees, and scripts used for phylogenetic analyses are available at Figshare (DOI: <u>10.6084/m9.figshare.31549345</u>).

### Septin structure predictions

To investigate potential interactions between *S. rosetta* septins, we used AlphaFold-3 (Abramson *et al*., 2024) to model the oligomeric septin assembly shown in Figure 1D and Figure S1B. As a point of comparison, we used the solved human septin hexamer cryo-EM structure (Mendonça *et al*., 2021; PDB ID: 7M6J). This hexamer comprises SEPT2G (NM_004404; a truncated form lacking N- and C-terminal domains to prevent higher-order polymerization), SEPT6 (NM_145799), and SEPT7 (AAH93640.2). The structure excludes unresolved regions of each septin, notably at the N- and C-termini of SEPT6 and SEPT7 (detailed information available at PDB and in File S1). No high-resolution structure of the canonical human septin octamer has yet been published, although its subunit order has been experimentally inferred (Kim *et al*., 2011), and inclusion of SEPT9 (NP_006631.1) in AlphaFold-3 predictions of the human octamer did not recapitulate this arrangement. Therefore, hexameric complexes were used for structural comparisons.

The *S. rosetta* septin hexamers in Figure 1D and Figure S1B were predicted using AlphaFold-3 (Abramson *et al*., 2024) with two copies each of *Sros_*SeptB (XP_004994452.1), *Sros_*Sept6 (XP_004992901.1), and *Sros_*SeptA (XP_004992637.1). The currently available RefSeq annotation for *Sros_*SeptB (XP_004994452.1) is truncated relative to the exon structure supported by RNA-seq read coverage from previous studies (Fairclough *et al*., 2013; Leon *et al*., 2025). We therefore derived a corrected gene structure using RNA-seq read coverage from Leon *et al*. (2025) for use throughout this study. The corrected CDS and protein sequences for *Sros*_*septB* are provided in File S2, and will be made available at GenBank under the accession number PZ120989 (pending final review). Because the PDB structure of the human septin hexamer omits unresolved residues at the N- and C-termini, we generated comparable *S. rosetta* protein sequences. To assign human–*S. rosetta* subunit pairs, we first modeled an untrimmed *S. rosetta* septin hexamer to define alignment pairs based on their relative positions within the hexamer (Figure S1B). We then aligned each *S. rosetta* septin to its paired, trimmed human counterpart using NCBI BLASTP and removed residues corresponding to the unresolved N- and C-terminal regions (File S1). These trimmed sequences were subsequently used for structural prediction.

The cryo-electron microscopy structure of the human septin hexamer was solved with guanosine diphosphate (GDP) bound to SEPT2 and SEPT7 subunits and guanosine triphosphate (GTP) bound to SEPT6 subunits (Mendonça *et al*., 2021). Accordingly, four GDP and two GTP molecules were included during AlphaFold-3 prediction of the *S. rosetta* hexamer. In the predicted model, GTP was associated with *Sros*_Sept6, whereas GDP was associated with *Sros*_SeptA and *Sros*_SeptB, consistent with subunit placement in the solved human structure.

Structures were visualized using UCSF ChimeraX v1.11.1 (Meng *et al*., 2023) for Figure 1D and for confidence visualization of the predicted *S. rosetta* hexamer shown in Figure S1B. Model confidence was assessed by Alphafold-3-generated predicted local distance difference test (pLDDT) scores and displayed as a continuous gradient (orange <50, yellow 50–70, cyan 70–90, blue >90), with 5-point transition zones on either side of cutoff values applied to smooth color boundaries. Global confidence metrics (pTM and ipTM) were obtained from AlphaFold-3.

### *S. rosetta* cell type RNA sequencing and analysis

To analyze septin expression across the life history of *S. rosetta* (Figure S2A), we used a recently published RNA-seq dataset (Leon *et al*., 2025). This dataset utilized updated cell state regulation protocols to analyze transcription across four populations: (1) slow swimmers and chains, (2) rosettes, (3) fast swimmers, and (4) thecates. Briefly, RNA-seq was performed in triplicate for each condition, and reads were mapped to the *S. rosetta* transcriptome using kallisto (Bray *et al*., 2016), with differential expression quantified using sleuth (Pimentel *et al*., 2017). Using this dataset, we compared expression of the four septin genes in *S. rosetta* between populations of slow swimmers and chains relative to the other three analyzed cell states. Average TPM values are presented in Figure S2A, and statistical comparisons are provided in the associated figure legend. Summary statistics are reported in Table S1. TPM values for individual replicates, fold-change, error metrics, and p- and q-values are given in File S3.

### Preparation of Rosette Inducing Factors released in outer membrane vesicles

In experiments where rosette induction was required, we used Rosette Inducing Factors released in outer membrane vesicles (RIF-OMVs) from the bacterium *Algoriphagus machipongonensis* (Alegado *et al*., 2013) (ATCC, BAA-2233). RIF-OMVs were provided as a gift from Florentine Rutaganira’s lab (Stanford, CA) and were prepared as described in Woznica *et al*. (2016) with a few modifications. A single colony of *A. machipongonensis* was isolated from an agar plate made with Seawater Complete (SWC) prepared in artificial seawater (ASW) as described in Levin and King (2013). The colony was used to inoculate 5 mL SWC medium and the culture was grown at 30°C with shaking. After 48 h, the 5 mL culture was transferred to 1 L of 100% SWC in a 2.8 L Fernbach flask and the culture was grown for 72 h at 30°C with shaking. After separating conditioned media from *A. machipongonensis* cells by spinning at 4000 × g for 30 min in 50 mL Falcon tubes and filtering the supernatant twice through a 0.22 µm filter, RIF-OMVs were isolated by centrifugation of the supernatant for 3 h at 37,000 rpm and 4°C in a Beckman Ti45 rotor. The membrane pellet was resuspended in a buffer of 50 mM HEPES-KOH, pH 7.4 and the solution of RIF-OMVs was filtered through a sterile 0.22 μm polyethersulfone syringe filter, aliquoted into 100 μL aliquots, flash frozen and lyophilized. Lyophilized RIF-OMVs were stored at −80°C. To assess induction efficiency, RIF-OMVs were resuspended in 100 µL DMSO and tested on *SrEpac* cultures across a dilution series (1:500–1:10,000), with dilutions of 1:500 to 1:5000 inducing rosettes at >70%. For this study, RIF-OMVs were instead resuspended in 100 µL 10 mM HEPES-KOH, pH 7.5. A final dilution of 1:1000, selected from the previously defined effective range, consistently induced robust rosette formation and was used across all rosette-induction experiments. Carrier controls consisted of an equivalent volume of 10 mM HEPES-KOH, pH 7.5.

### Choanoflagellate culturing

All experiments were performed using strains of *S. rosetta* co-cultured with *Echinicola pacifica* bacteria (strain designation: *SrEpac*) (Levin and King, 2013). Throughout this study, cells were grown in low nutrient media (LNM) (Booth and King, 2020). LNM consists of AK sea water (AKSW) complete (AKSWC; 5.0 g/L peptone, 3.0 g/L yeast extract, and 3.78 g/L glycerol dissolved in AK seawater) and AK cereal grass media (AKCGM3; 5 g/L cereal grass steeped in AKSW (Carolina Biological Supply Company, Cat. No. 132375)), both diluted to 1.5% by volume in AKSW and supplemented with nitrates, phosphates, trace elements, and vitamins as specified in Booth and King (2020). The AKSW formulation has been used to culture marine algae (Hallegraeff *et al*., 2004) and dinoflagellates (Skelton *et al*., 2009) and was adapted for culturing *S. rosetta* in Booth *et al*. (2018) because its lower calcium concentration (2.7 mM) was hypothesized to be more suitable for transfection protocols in comparison to the routinely used ASW made from Tropic Marin sea salts (Tropic Marin) (Levin and King, 2013). AKSW has since become one of the standard seawater options for *S. rosetta* husbandry.

A modified media formulation, comprised of AKSWC and AKCGM3 diluted each to 4% by volume in AKSW, termed high nutrient media (HNM), permits higher maximum *S. rosetta* cell densities during log-phase culture (Booth and King, 2020) and has been used in recent studies (Booth and King, 2020; Coyle *et al*., 2023). We instead used LNM as it suppressed *E. pacifica* biofilm formation during routine passaging while supporting cell densities sufficient for experimental use.

Cultures were passaged every two days at a 1:500 dilution into fresh LNM and supplemented with 0.4% (v/v) of a 10 mg/mL *E. pacifica* suspension in ASW (hereafter referred to as “*E. pacifica* food pellet”) to ensure a consistent supply of food bacteria. Cultures were maintained at 22°C and 60% humidity. For consistency, all experiments were performed using cells in the mid-log phase of growth, which in this media formulation occurs between 1 × 10^5^ and 1 × 10^6^ cells/mL.

*S. rosetta* cells grown to log-phase without RIF-OMVs produce cultures containing both single-celled slow swimmers and chain colonies of varying sizes (Figure 2A). When retention of chain colonies was not required, routine handling steps often disrupted chains, resulting in cultures composed primarily of single cells (Figure 3A; Figure S3B). In contrast, growth in the presence of RIF-OMVs produced cultures containing cells at multiple stages of rosette development. Rosettes are arbitrarily defined as starting at the four-cell stage (Larson *et al*., 2020), although they start as single cells and develop through one-, two-, and three-cell stages en route to becoming rosettes. Therefore, depending on the time since induction or the application of shear, cultures also contained different percentages of single cells, doublets, and triplets (Figure S4G). Induction for at least 24 h resulted in robust rosette formation in wild-type cells, with colonies of approximately 8 – 12 cells predominating in the resulting cultures.

For long-term storage of wild-type and mutant *SrEpac* strains, cryopreservation was performed using 10% DMSO as described in King *et al*. (2009). Midway through this study, a glycerol-based cryopreservation method was reported and is now preferred (Chandra and Rutaganira, 2025); however, the DMSO protocol was used throughout our study for consistency across strains.

### CRISPR guide RNA and repair template design

CRISPR guide RNA (gRNA) and repair templates were designed following previously established protocols (Booth and King, 2020). Candidate CRISPR RNA (crRNA) sequences for each of the four septin genes were identified using the EuPaGDT tool (http://grna.ctegd.uga.edu/) and the *S. rosetta* genome. Guide length was set to 15 nucleotides and an expanded PAM consensus sequence (HNNRRVGGH) was used. Coding sequences for genes of interest were obtained from the Ensembl Protists database (Yates *et al*., 2022), which hosts the *S. rosetta* genome, with the exception of the corrected coding sequence for *Sros_septB* as described in “Septin structure prediction” (File S2). crRNA candidates were filtered for guides with one on-target hit (including making sure the guides do not span exon-exon boundaries), zero off-target hits (including against the genome of the co-cultured bacterium *E. pacifica*), lowest strength of the predicted secondary structure (assessed using the RNAfold web server: http://rna.tbi.univie.ac.at/cgi-bin/RNAWebSuite/RNAfold.cgi), and proximity to the 5’ end of the coding sequence to maximize early gene disruption. crRNAs with the guide sequence of interest, as well as universal trans-activating CRISPR RNAs (tracrRNAs), were ordered from IDT (Integrated DNA Technologies) (File S4).

Repair templates were designed as single-stranded DNA oligonucleotides matching the sense strand of the crRNA and containing homology arms corresponding to 50 bp of genomic sequence flanking each side of the Cas9-induced double-stranded break at the CRISPR target site. For gene truncation experiments, the sequence 5’-TTTATTTAATTAAATAAA-3’, a “termination cassette” encoding a stop codon in every frame, was included between the homology arms. To ALFA-tag (Götzke *et al*., 2019) *Sros*_*septA,* a codon-optimized ALFA-tag sequence (5’-AGCCGCCTGGAGGAGGAGCTGCGCCGCCGCCTGACGGAG -3’) was inserted between homology arms. A 5’ AG and 3’ G were included to restore the interrupted GAG (glutamic acid) codon at the cut site (pos. 1178 in the CDS sequence) (File S4). Along with the crRNA targeting our gene of interest, we co-edited our cells using previously published sequences for a crRNA and repair template designed to introduce a P56Q mutation in *rpl36a* (Booth and King, 2020), conferring cycloheximide resistance to enrich for genome-edited cells by selection. crRNA and repair oligo sequences are given in File S4.

### Genome editing

Thirty-six hours prior to transfection, *S. rosetta* cells were inoculated into 120 mL of LNM at 8,000 cells/mL with 1 mL of *E. pacifica* food pellet added. This seeding density brought the culture to mid-log phase at the time of transfection. The inoculation period and bacterial food pellet addition were adjusted from previous protocols (Booth and King, 2020) to better match growth dynamics in LNM.

CRISPR/Cas9 nucleofection was performed according to the protocol outlined in Coyle *et al*. (2023), which adapted the original protocol reported in Booth and King (2020). Cultures were co-transfected with Cas9 RNPs and repair oligonucleotides targeting (1) the gene of interest and (2) the *rpl36a* locus to confer cycloheximide resistance (see “CRISPR guide RNA and repair template design”). The nucleofection reactions were prepared as described in Coyle *et al*. (2023), mixed into Lonza SF buffer (Lonza, Cat. No. V4SC-2960), added to a 96-well nucleofection plate (Lonza, Cat. No. V4SC-2960), and pulsed with the program CM156 using a Lonza 4D-Nucleofector (Cat. No. AAF-1003B for the core unit and AAF-1003S for the 96-well unit). Pulsed cells were immediately recovered in recovery buffer (HEPES-KOH, pH 7.5; 0.9 M sorbitol; 8% [wt/vol] PEG 800) for 5 min, and then added to 2 mL LNM in a 6-well plate (Thermo Fisher Scientific, Cat. No. 140675) at 22°C and 60% humidity. After 30 min, 10 μL *E. pacifica* food pellet was added. Twenty-four hours later, 10 μl of 1 μg/mL cycloheximide (final concentration 5 ng/mL) was added to each well along with an additional 10 μl *E. pacifica* food pellet. Cycloheximide selection was carried out for four days. A negative control well of unedited cells (carried through the transfection experiment but without Cas9 RNPs or repair oligonucleotide) was used to confirm effective cycloheximide selection after the four-day selection period.

Following selection, cells were diluted to 100,000 cells/mL and clonally isolated using a WOLF® G2 Cell Sorter (WOLF; NanoCellect Biomedical) equipped with a WOLF Microfluidic Sorting Cartridge (Cat. No. SP0511). Cells were sorted into 96-well plates (Thermo Fisher Scientific, Cat. No. 167008) containing 200 μL LNM supplemented with 1 μL of *E. pacifica* food pellet per well. Sorting was performed using forward scatter as the trigger, with a threshold of 40,000 and detector voltage set to 150. These conditions consistently produced plates with cells in ∼75% of the wells. For each independent population of edited cells, at least four 96-well plates were prepared. Sorted cells were allowed to proliferate for five days to reach workable cell density within wells.

To extract genomic DNA for genotyping analysis, 25 μl of culture from the same well position in up to four source plates (e.g., A1 from each plate) was pooled into the corresponding well of a new 96-well plate. Pooling was performed to reduce the number of sequencing reactions required for genotyping large numbers of cells. An equal volume of DNAzol direct (Molecular Research Center, Inc, Cat. No. DN131) was added to the pooled cultures to extract genomic DNA. This mixture was incubated at room temperature for 10 min and then stored at -20°C.

Genotyping PCRs were performed in 96-well plates (Brooks Life Sciences, Cat. No. 4ti-0770/c) using Q5 polymerase (NEB, Cat. No. M0491L) for 40 cycles. 2.5 μl of genomic DNA template was used in each 25 μl PCR reaction. PCR products were purified by magnetic bead cleanup and analyzed by Sanger sequencing (UC Berkeley DNA Sequencing Facility). Positive edits were detected in pooled genomic DNA samples by identifying overlapping sequencing reads beginning immediately downstream of the designed edit site. After a candidate positive pool was identified, the corresponding wells from the original 96-well plates were sequenced individually to identify the positive well, and the resulting strain was expanded for downstream experimentation and long-term storage.

### Development of CRISPR/Cas9-mediated endogenous C-terminal fluorescent tagging in *S. rosetta*

To visualize *Sros*_SeptA in live cells, we adapted a recently published CRISPR/Cas9 gene truncation strategy (Combredet *et al*., 2025) to enable endogenous fluorescent tagging in *S. rosetta*. Our approach enabled in-frame insertion of the coding sequence of mStayGold (Ivorra-Molla *et al*., 2024), a bright and photobleach-resistant monomeric fluorophore, at the 3’ end of the *Sros*_*septA* coding sequence to generate an endogenously tagged C-terminal fusion protein *Sros*_SeptA-mStayGold.

The Combredet *et al*. (2025) method uses a plasmid template (pEFL-pac-5’ Act; Addgene ID pMS18; Cat. No. 225681) to PCR-amplify a repair template for use in CRISPR/Cas9 genome editing. In the published strategy, the repair template contains (1) a backbone-derived puromycin resistance cassette (pEFL-pac-5’ Act), (2) a primer-derived termination cassette encoding a premature stop codon, and (3) primer-derived 5’ and 3’ homology arms corresponding to the genomic sequence flanking each side of the Cas9-induced double-stranded break at the CRISPR target site. Combredet *et al*. tested 50, 80, and 155 bp homology arms, all of which resulted in similar editing efficiency. Following CRISPR/Cas9-mediated double-strand break formation, the repair template is inserted at the target locus, resulting in gene truncation and enabling puromycin selection of edited cells.

We devised a similar strategy to tag *Sros*_SeptA at its C-terminus with a fluorescent protein. First, we identified the most 3’-adjacent CRISPR/Cas9 PAM site within the *Sros*_*septA* coding sequence. We then modified the pMS18 plasmid to encode: (1) the 59 bp region of the *Sros*_*septA* CDS downstream of the cut site (“*Sros*_*septA* 3’ CDS recovery region”), (2) a glycine-serine flexible linker sequence (SGGSGGS) (Chen *et al*., 2013), fused in frame, which had previously been used to link *Sros*_SeptA to an mWasabi fluorophore (Booth *et al*., 2018), (3) an in-frame *mStayGold* coding sequence ending with a stop codon, and (4) the 3’ UTR of *Sros*_*septA*, all positioned immediately upstream of the puromycin-resistance cassette. The *mStayGold* coding sequence, codon-optimized for *S. rosetta*, was generously provided by Jeffrey Colgren (Pawel Burkhardt lab, University of Bergen). The resulting plasmid is available at Addgene (Addgene ID NK812; Cat. No. 253995). We designed PCR primers to amplify the modified repair cassette (*Sros*_*septA* 3’ CDS recovery region – 21 bp linker – *mStayGold* coding sequence – stop codon – puromycin resistance cassette) with 50 bp homology arms flanking the CRISPR/Cas9 target site in the *Sros*_*septA* coding sequence, generating a repair template size 2996 bp. Detailed information about the sequences described above are available in File S5.

To avoid possible recombination between two identical 50 bp sequences on the repair template – (1) the homology arm sequence of the 3’ primer, designed to match the genomic region adjacent to the cut site, and (2) the 3’ *Sros*_*septA* CDS recovery sequence encoded on the plasmid template – we introduced eight synonymous nucleotide substitutions into the plasmid-encoded CDS recovery region to reduce similarity between these sequences (File S5). All substitutions were placed at third codon positions such that the edited genomic allele encodes an unchanged *Sros*_SeptA amino acid sequence. The alternative codons were selected to match the original codon usage as closely as possible (File S5).

Following PCR amplification and cleanup of the 2996 bp repair template as outlined in Combredet *et al*. (2025), we recovered ∼9 μg of repair template contained in 30 μL nuclease-free water. We then evaporated the 30 μL volume containing the repair template at 55°C on a heating block for 3 h until near dryness and resuspended the repair oligos directly in the final CRISPR reaction mixture (16 μL ice-cold “homebrew” nucleofection buffer (Nguyen *et al*., 2026), 4 μL SpCas9 RNP, 1 μg pUC19 (Nature Technology, Cat. No. NTC-RP500) carrier plasmid, and 2 μL of primed *S. rosetta* cells. SpCas9 RNP and primed *S. rosetta* cells were prepared as in Booth and King (2020) with the modifications given in “Genome editing”. The nucleofection reaction was then added to a 16-well Lonza nucleofection strip (Lonza, Cat. No. V4XP-3032) and pulsed with the program DG137 (Nguyen *et al*., 2026) using a Lonza 4D-Nucleofector. Pulsed cells were immediately recovered in recovery buffer for 5 min and then added to 2 mL LNM in a 6-well plate. After 30 min, 10 μL *E. pacifica* food pellet was added. Twenty-four hours after nucleofection, 80 μg/mL puromycin was added to the culture and selection took place over five days. Healthy cells that recovered following puromycin selection were then screened for fluorescence on a Nikon Ti2 inverted microscope (Nikon Instruments) with a Yokogawa CSU-W1 spinning disk confocal unit and a Hamamatsu ORCA-Flash4.0 C14440-20UP CMOS camera (Hamamatsu Photonics) (hereafter, “Nikon Ti2 spinning disk confocal system”). Fluorescent cultures were then clonally isolated and genotyped by PCR and Sanger sequencing as described in “Genome editing” (primers listed in File S5).

### Growth curves

Twenty-four hours before the start of the growth assay (Figure S2C), cultures in log-phase were diluted to 10,000 cells/mL to standardize the growth conditions across starter cultures. On the day the growth curve plates were seeded, starter cultures were washed three times (2,400 × g for 5 min each) with ASW to remove bacteria on a Thermo Fischer Scientific ST16R swinging-bucket rotor centrifuge (hereafter, “swinging-bucket rotor centrifuge”), thereby standardizing nutrient conditions across cultures. Cells were then diluted to 5,000 cells/mL in LNM and supplemented with 267 µg/mL *E. pacifica* food pellet. 500 μL of culture was aliquoted into each well of a 24-well plate (Fisher Scientific, Cat. No. 09-761-146) and incubated at 22°C and 60% humidity. Plates were maintained in a covered plastic container with dampened paper towels and the lid loosely affixed to minimize evaporation while allowing gas exchange.

Every 12 h, starting at 0 h and continuing for 96 h, three wells per strain were fixed with 10 μL of 16% paraformaldehyde (Thermo Fisher Scientific, Cat. No. 50-980-487) and stored at 4°C. After all time points were collected, samples were vortexed, and 200 μL of each was added to a 96-well plate (Thermo Fischer Scientific, Cat. No. 167008). Samples were analyzed on an Attune CytPix Flow Cytometer equipped with an Attune CytKick Max Autosampler (Thermo Fisher Scientific) with the following settings: Mix Mode (aspirate and stir once per well); 150 µL acquisition volume at a flow rate of 200 µL/min; FSC detector voltage 150; SSC detector voltage 300; FSC threshold 20 × 10³. Total cell counts were divided by the acquisition volume to calculate cell density (cells/mL).

### Imaging slow swimmers, chains, and rosettes

To image *S. rosetta* wild-type and septin mutant cultures shown in Figure 2B–F’’, Figure S2D, and Figure S4D, cultures were grown for 36 h in six-well plates (Thermo Fisher Scientific, Cat. No. 140675) (3 mL per well) with or without RIF-OMVs. For RIF-OMV-induced cultures subjected to shear, cultures were vortexed for 15 s at high-speed using a Vortex Genie 2 (Scientific Industries) in a 15 mL Falcon tube (Corning, Cat. No. 352196) before imaging.

For each strain and condition, 2 mL of culture was gently concentrated by low-speed centrifugation (600 × g for 5 min) using a swinging-bucket rotor centrifuge. To avoid disturbing the loose pellet, the supernatant was left in place and a 200 μL aliquot was gently aspirated directly from the pellet using a wide-bore 200 μL tip and transferred to a poly-D-lysine (Millipore Sigma, Cat. No. P6407) coated 8-well FluoroDish (World Precision Instruments, Cat. No. FD35-100). The gentle centrifugation and transfer preserved chain and rosette structures while concentrating cells for ease of imaging. Cells were allowed to settle for 10 min at 22°C before live imaging by differential interference contrast (DIC) microscopy on a Zeiss Axio Observer.Z1/7 Widefield microscope with a Hamamatsu Orca-Flash 4.0 LT CMOS Digital Camera (Hamamatsu Photonics) (hereafter, “Zeiss Axio Observer Widefield microscope”) equipped with a Plan-Apochromat 20×/0.80 Ph 2 M27 objective with a 1.6× Optovar setting.

### Imaging and quantifying cell area of slow swimmers and chain cells

To quantify the cell area of slow swimmers and chain cells (Figure 2G; Figure S2E and S2F), cultures were grown to mid-log phase and visually inspected prior to analysis to confirm that only slow swimmers and chains, and not smaller fast swimmer cells, were present. This was done to ensure cell size comparisons were consistent across strains. Cells were then vortexed at high speed for 5 s to separate doublets and chain colonies into single cells before analysis.

Cells were analyzed on an Attune CytPix Flow Cytometer, which captures brightfield images of recorded events in addition to standard flow cytometry data, using the following settings: 100 μL acquisition volume at a flow rate of 200 μL /min; FSC detector voltage 150; SSC detector voltage 300; FSC threshold 30 × 10^3^; and 10,000 images captured total. The “Cells_Full_Resolution_v23” stock CytPix image processing model (available at Figshare, DOI: <u>10.6084/m9.figshare.31549345</u>) was used to process images and determine the projected area of each cell. Only images containing a single cell were included in the final data set. When the model detected two or more cells in an image (fewer than 5% of images analyzed), those images were excluded (Figure S2E).

### Imaging and quantifying rosette assembly and integrity phenotypes

To quantify the projected area of individuals (single cells, doublets, triplets, and rosettes) in rosette-induced cultures with and without exposure to shear (Figure 2H), starter cultures in mid-log phase were split and used to seed duplicate 2 mL volumes in six-well plates. Cultures were then grown for 24 h in the presence of RIF-OMVs. To apply shear treatment, cultures were vortexed for 15 s at high speed in a 15 mL Falcon tube using a Vortex Genie 2. Vortexed cultures were allowed to rest for 10 min prior to subsequent preparation steps to minimize potential cell lysis. For each condition, 1 mL each of vortexed and unvortexed culture were gently transferred to a FluoroDish using a 1 mL pipette tip with the end trimmed off. Cells were allowed to settle for 10 min prior to staining.

LysoTracker Red DND-99 (Thermo Fisher Scientific, Cat. No. L7528) was added at a final dilution of 1:100 (overloaded to visualize the cell body) by removing 500 μL of culture from the FluoroDish and carefully adding 500 μL of a 1:50 LysoTracker solution diluted in ASW. Cells were immediately imaged using DIC microscopy and epifluorescence on a Zeiss Axio Observer Widefield microscope equipped with a Plan-Apochromat 20×/0.80 Ph 2 M27 objective with a 1.6× Optovar setting. Epifluorescence images were acquired using a mCherry filter set. At least five fields of view were acquired per condition for each biological replicate. Fields of view were chosen through sequential non-overlapping stage translations; fields with insufficient cell density or poor staining were excluded.

Images were batch processed in FIJI (ImageJ) (Schindelin *et al*., 2012) to ensure consistent analysis across samples. The epifluorescence channel of each image was converted to 8-bit and thresholded using the Otsu method (upper limit set to 150) to create a binary mask; the mask was inverted so that objects were black on a white background prior to particle analysis; segmented objects were analyzed with the “Analyze Particles” command to calculate the area of each individual; and a minimum size threshold of 10 pixels^2^ was set to exclude small artifacts. Examples of DIC images, epifluorescence images, and masked images are given in Figure S2G. The FIJI batch protocol is available at Figshare (DOI: <u>10.6084/m9.figshare.31549345</u>).

### Imaging and quantifying cell volume of slow swimmers, chains, and rosettes

To quantify the average cell volume of cells in rosettes in comparison with slow swimmer cells and cells in chains (Figure 3B), cultures were grown for 36 h in six-well plates (3 mL per well) with or without RIF-OMVs, respectively. 1 mL of each culture was added to a poly-D-lysine-coated FluoroDish and stained with LysoTracker Red DND-99 as in “Imaging and quantifying rosette assembly and integrity phenotypes.” Cells were then immediately imaged using a Nikon Ti2 spinning disk confocal system equipped with an Apo TIRF 60×/1.49 NA oil immersion objective. Fluorescent images were acquired using 561 nm excitation with a mCherry emission filter. Three-dimensional imaging was performed with a 0.2 µm step size over ∼90 optical sections (total depth ∼18 µm). At least four fields of view were acquired per condition for each biological replicate. Fields of view were chosen through sequential non-overlapping stage translations; fields with insufficient cell density or poor staining were excluded.

Images were batch processed to quantify cell volume using a cell segmentation pipeline within Imaris x64 v10.2.0 (Jun 18, 2024 build 67164, Bitplane/Oxford Instruments) to ensure consistency across strains (example images before and after segmentation shown in Figure S3B). To segment single cells and cells in rosettes, surfaces were generated from the raw, unprocessed three-dimensional images using the machine learning–based segmentation workflow option in Imaris by iterative manual annotation of foreground and background objects. To train the segmentation model, images of representative untreated and rosette-induced cells from each genotype (wild-type, *Sros*_*septA^1-50^*, and *Sros*_*septA^1-116^*) were combined into a composite training image. The final model was trained by annotating at least five cells from each genotype and treatment condition within this composite image. Surface generation was performed using the region growing segmentation method with an estimated object diameter of 1.5 µm and surface smoothing enabled (surface grain size = 0.216 µm). Following segmentation, objects were filtered to exclude surfaces with fewer than 10 voxels, volumes less than 20.0 µm³, and objects intersecting the image border in the XY plane (segmented example images given in Figure 3A). A batch protocol was created from these parameters and applied to images in all conditions, resulting in cell volume as an output. The training data and segmentation parameters used in Imaris are available at Figshare (DOI: <u>10.6084/m9.figshare.31549345</u>).

### Imaging and quantifying cytokinesis failure in slow swimmers and rosettes

For imaging (Figure 3C) and quantification (Figure 3D) of cytokinesis events, wild-type and *Sros*_*septA^1-50^* cells were diluted to 150,000 cells/mL and gently vortexed for 5 s on low speed on a Vortex Genie 2 to separate chains and doublets into single cells. 200 μL of each diluted culture with or without RIF-OMVs and supplemented with 2 μL *E. pacifica* food pellet was added to the center of a FluoroDish. Dishes were prepared by coating with poly-D-lysine, washing three times with ASW, and leaving the final wash in place for 30 min to generate a gently adhesive surface that minimized surface effects on cell division. To minimize evaporation while permitting oxygen exchange during time-lapse imaging, 1.5 mL of Anti-Evaporation Oil (Ibidi, Cat. No. 50051) was layered on top of the 200 μL culture, enough to cover the FluoroDish surface and with an oil level matching the height of the culture droplet. Cells were then allowed to settle for 10 min at 22 °C.

Cells were imaged using DIC time-lapse microscopy on a Zeiss Axio Observer Widefield microscope equipped with a Plan-Apochromat 20×/0.80 Ph2 M27 objective. For each experiment, a single field of view with a high density of cells was chosen. Images were collected at 30 s intervals for 10 h. A cytokinesis failure was defined as a cell division event followed by cell fusion at any point within 30 min after the initial event of cleavage furrow ingression could be detected. All division events across each time-lapse were recorded and included in quantification, except for divisions beginning within the last 30 min of the time-lapse as they could not be analyzed with the above parameters.

### Imaging *Sros*_SeptA-mStayGold

To image *Sros*_SeptA-mStayGold slow swimmers, chains, and rosettes (Figure 4B), cultures were grown for 12 h in six-well plates (3 mL per well) with or without RIF-OMVs. This induction time was optimized to enrich for chains and rosettes at the four-cell stage for ease of imaging. 2 mL of each culture was first gently concentrated by low-speed centrifugation (600 × g for 5 min). To avoid disturbing the loose pellet, the supernatant was left in place and a 200 μL aliquot was gently aspirated directly from the pellet using a wide-bore 200 μL tip. The aspirated pellet was then transferred to a poly-D-lysine-coated 96-well plate to preserve chain and rosette structures. *Sros*_*septA-mStayGold* cultures were stained with Alexa Fluor 568-conjugated Jacalin (Vector Labs, Cat. No. L-1150; Thermo Fischer Scientific, Cat. No. A10238) at 1:200 and imaged immediately using a Nikon Ti2 spinning disk confocal system operating in a 2× super-resolution (SoRa) mode with a Plan Apo λ 100× oil immersion objective. Fluorescent images were collected using 488 nm excitation with a FITC emission filter (SeptA-mStayGold) and 561 nm excitation with a TRITC emission filter (Jacalin). Z-stacks were collected at 0.2 µm intervals and representative central optical sections are shown in Figure 4B.

For time-lapse imaging to visualize SeptA-mStayGold during cell division (Figure 4C and S4H and Videos S4 and S5), *Sros*_*SeptA-mStayGold* cells were grown for 6 h in six-well plates (3 mL per well) with or without RIF-OMVs. This induction period was optimized to enrich for early rosette development for ease of time-lapse, three-dimensional imaging. Various lectins, including Jacalin and WGA, were tested to label dividing cells; however, lectins appeared to inhibit cell division and were therefore not used for time-lapse imaging. 2 mL of culture was gently concentrated by low-speed centrifugation (600 × g for 5 min) with a swinging-bucket rotor centrifuge. A 200 μL aliquot was carefully collected from the resulting loose pellet using a wide-bore 200 μL pipette tip, transferred to a 96-well plate, and imaged using the same microscope system and excitation/filter set as above. Z-stacks were acquired at 0.2 µm intervals and time points were acquired at 30 s intervals. The three-dimensional time-lapse images were visualized and processed using Imaris x64 v10.2.0.

### Immunostaining and imaging *Sros*_SeptA-ALFA

To visualize the cellular localization of SeptA-ALFA in interphase cells (Figure S4B), 20 mL of culture was grown for 12 h with or without RIF-OMVs. Cells (13 mL) were collected by centrifugation (1000 × g for 5 min) in 15-mL Falcon tubes using a swinging-bucket rotor centrifuge and resuspended in LNM to a cell density of 5 × 10^6^ cells/mL (∼1 mL). Cells were fixed in suspension prior to plating: A 200 μL aliquot of each culture was pre-treated in an Eppendorf tube in 6% acetone in phosphate-buffered saline (PBS) at room temperature for 5 min followed by fixation in 4% PFA in PBS at room temperature for 10 min. Fixed cells were spun for 10 min at 800 × g with an Eppendorf 542R fixed-angle tabletop microcentrifuge and transferred to room temperature PEM (100 mM PIPES, pH 6.95; 2 mM EGTA; 1 mM MgCl^2^). At this point, cells could be stored at 4 °C until immunostaining steps.

To prepare for immunostaining, cells in PEM were transferred to a poly-D-lysine–coated plate and centrifuged (600 × g for 5 min) with a swinging-bucket rotor centrifuge to adhere fixed cells. Cells were permeabilized with permeabilization buffer (1% [w/v] bovine serum albumin-fraction V; 0.3% [v/v] Triton X-100 in PEM) for 30 min. Cells were then incubated with 12.5 nM anti-ALFA nanobody conjugated to Alexa Fluor 657 (NanoTag Biotechnologies, Cat. No. N1502- AF647-L) and 25 ng/mL rat anti-Tubulin antibody conjugated to Alexa Fluor 488 (Abcam, Cat. No. Ab195883) for 1 h while rocking at 300 rpm on an Eppendorf MixMate (Eppendorf, Cat. No. 5353953449).

Chambers were then washed three times with PEM and imaged immediately using a Nikon Ti2 spinning disk confocal system equipped with a Plan Apo λ 100× oil immersion objective. Fluorescent images were collected using 488 nm excitation with a FITC emission filter (microtubules) and 647 nm excitation with a Cy5 emission filter (SeptA-ALFA). Z-stacks were acquired at 0.3 µm intervals. The three-dimensional fluorescent images were then visualized and processed using Imaris x64 v10.2.0.

To visualize SeptA-ALFA-positive intercellular bridges (Figure S4C), 20 mL of uninduced culture was grown to early log-phase, which we found contained the highest amount of chain colonies. 13 mL of culture was gently added to a 15 mL Falcon tube and concentrated by low-speed centrifugation (600 × g for 5 min) with a swinging-bucket rotor centrifuge. To avoid disturbing the loose pellet, 10 mL of supernatant was removed and a 200 μL aliquot was then gently aspirated directly from the pellet using a wide-bore 200 μL tip. The aspirated pellet was then transferred to a poly-D-lysine-coated 96-well plate to preserve chain and intercellular bridge structures. One minute before fixation, 1 μg/mL FM-4-64FX dye (Thermo Fischer, Cat. No. F34653) was added to the cultures to label membranes. Cells were then fixed directly in the chamber using 6% acetone, 4% PFA, and 1:300 phalloidin conjugated to Alexa Fluor 488 (Life Technologies, Cat. No. A12379) in PBS at room temperature for 15 min in the dark. Acetone, PFA, and phalloidin were added simultaneously as we noticed this combination, along with gentle culture handling, preserved a higher number of intercellular bridges in chain colonies compared with previously established fixation protocols (Booth *et al*., 2018), potentially due to phalloidin stabilizing intercellular bridge structures, which often contained F-actin. Following fixation, wells were washed twice with PEM and then incubated in permeabilization buffer as above on a shaker at 300 rpm for 30 min. Cultures were then incubated with 12.5 nM anti-ALFA nanobody conjugated to Alexa Fluor 657 on a shaker at 300 rpm for 1 h. Cultures were washed three times with PEM and imaged immediately as above with the following changes: Z-stacks were acquired at 0.2 µm intervals, 488 nm excitation with a FITC emission filter was used to visualize phalloidin-labeled F-actin, and 488 nm excitation with a Cy5 emission filter was used to visualize FM4-64-labeled membrane. Maximum-intensity projections were generated from a total Z-depth of 1 μm.

### Imaging the extracellular matrix of rosettes

To visualize Jacalin-positive ECM in wild-type and *Sros*_*septA^1-50^* (Figure S3A), cultures were grown for 12 h in six-well plates (3 mL per well) with or without RIF-OMVs. 200 μL of culture was then added to poly-D-lysine–coated 96-well plates to allow cells to adhere, and fluorescein-conjugated Jacalin was added at a final dilution of 1:200 by removing 100 μL culture and adding 100 μL 1:100 fluorescein-conjugated Jacalin premixed in LNM. Cells were immediately imaged by DIC and epifluorescence microscopy using a Zeiss Axio Observer Widefield microscope equipped with a LD C-Apochromat 40×/1.1 W Korr UV VIS IR water-immersion objective with a 2.6× Optovar setting. Epifluorescence images were acquired using a GFP filter set. Z-stacks were collected at 1 μm step sizes. Maximum-intensity projections were generated from a total Z-depth of 5 μm for the fluorescence channel, and the corresponding central DIC plane was used for merged images.

To visualize Rosetteless-positive ECM in wild-type and *Sros*_*septA^1-50^* (Figure S3A’), 20 mL cultures were grown to mid-log phase after 24 h of induction with RIF-OMVs. Rosetteless, a C-type lectin required for rosette formation in *S. rosetta*, has previously been shown to localize to rosette ECM (Levin *et al*., 2014). Cells were concentrated, fixed, washed, and transferred to 96-well plates as described in the first section of “Immunostaining and imaging SeptA-ALFA.” Cells were permeabilized with permeabilization buffer (1% [w/v] bovine serum albumin-fraction V; 0.3% [v/v] Triton X-100 in PEM) for 30 min and then stained with 1:200 rabbit anti-Rosetteless (Levin *et al*., 2014) in blocking buffer (1% [w/v] bovine serum albumin-fraction in PEM). Cells were incubated with the primary antibody for 1 h while rocking at 300 rpm on an Eppendorf MixMate. The chamber was then gently washed three times with PEM and incubated with a final concentration of 2 µg/mL goat anti-rabbit secondary antibody conjugated to Alexa Fluor 568 (Thermo Fisher Scientific, Cat. No. A-11036) for 45 min while rocking at 300 rpm.

Following incubation with the secondary antibody, the chamber was washed three times with PEM and cells were imaged immediately using a Zeiss Axio Observer Widefield microscope equipped with a LD C-Apochromat 100×/1.46 Oil DIC (UV) M27 objective with a 1.6× Optovar setting. Epifluorescence images were acquired using a Cy3 filter set. Z-stacks were collected at 1 μm step sizes. Maximum-intensity projections were generated from a total Z-depth of 5 μm for the fluorescence channel, and the corresponding central DIC plane was used for merged images.

### Phenotypic comparison of tagged *Sros*_*septA* strains

To compare phenotypes of wild-type, *Sros*_*septA^1-50^, Sros*_*septA-ALFA^392^,* and *Sros*_*septA-mSG* rosettes, cultures were grown for 24 h in six-well plates (3 mL per well) with RIF-OMVs. Cultures were prepared in duplicate and experiments to quantify cell volume (Figure S4E) and rosette integrity (Figure S4F) were performed on the same days. On one of the days, cultures were prepared in triplicate and representative DIC images (Figure S4D) images were also captured. To compare cell volume within wild-type, *Sros*_*septA^1-50^, Sros*_*septA-ALFA^392^,* and *Sros*_*septA-mSG* rosettes, microscopy and cell segmentation were performed as described in “Imaging and quantifying cell area and volume.” The shorter rosette induction period used here may have contributed to the reduced severity of the *Sros*_*septA^1-50^* cell size phenotype observed in Figure S4E compared to Figure 3B.

To analyze the rosette development phenotypes in wild-type, *Sros*_*septA^1-50^, Sros*_*septA-ALFA^392^,* and *Sros*_*septA-mSG*, we developed an imaging pipeline capable of resolving the number of cells per individual for single cells, doublets, triplets, and rosettes using the Attune CytPix Flow Cytometer (Figure S4G). Using this pipeline, we analyzed septin-tagged rosettes after application of shear force to quantify rosette development (i.e. the combined phenotypes of rosette assembly and rosette integrity) in a single, high-throughput metric. Rosette-induced cultures were first exposed to shear through vortexing at high speed for 15 s using a Vortex Genie 2. The resulting cultures were analyzed on the Attune CytPix using the following settings: 400 µL acquisition volume at a flow rate of 200 µL/min; FSC detector voltage 150; SSC detector voltage 300; FSC threshold 30 × 10³; 5,000 images acquired. To mask the area of individuals and then identify number of cells within those individuals, we trained a custom segmentation and cell identification pipeline by iterative manual annotation of foreground and background objects until cell surfaces were accurately segmented. The custom pipeline was trained on thirty test images from across (1) the four strains analyzed and (2) a diversity of multicellular morphs to create a representative training model. Our CytPix user-trained model is available at Figshare (DOI: 10.6084/m9.figshare.31549345).

This pipeline was not used to analyze the rosette assembly and integrity defects of septin mutants in Figure 2H because, although the CytPix is gentle, it may still disrupt rosettes in non-vortexed cultures and thereby confound measurements of rosette assembly. Conversely, the analysis pipeline used for Figure 2H was not applied to the tagged septin strain comparison in Figure S4F because we instead combined the rosette assembly and integrity phenotypes into a single metric that avoided measurement bias arising from differences in cell size. In Figure 2H, cell size bias was present in between-strain comparisons of rosette assembly; however, phenotypic differences were sufficiently large that this effect was considered acceptable. By contrast, rosette integrity comparisons done in Figure 2H were performed within strains, minimizing cell size bias for this metric.

For the representative images of wild-type, *Sros*_*septA^1-50^, Sros*_*septA-ALFA^392^,* and *Sros*_*septA-mSG* rosettes before and after vortexing (Figure S4D), rosettes were either untreated or vortexed at high speed for 15 s as described above and imaged as described in “Imaging single cells, chains, and rosettes.”

### Statistics

Information about the quantification and statistical details of experiments can be found in the corresponding figure legends. Statistical tests and graphs were produced using Prism 9.0.0.

## DATA, SCRIPT, REAGENT AND STRAIN AVAILABILITY

Plasmids in this study have been deposited to Addgene (Addgene ID NK812; Cat. No. 253995). Choanoflagellate cell lines used in this study are available upon request. Sequence datasets, multiple sequence alignments, phylogenetic trees, and image-analysis scripts used in this study are available at Figshare (DOI: 10.6084/m9.figshare.31549345).

## ACKNOWLEDGMENTS

We thank the following people for insights and contributions that helped advance this work: Maxwell Coyle, Juliette Mathiue, and Agathe Chaigne provided valuable feedback on drafts of the manuscript. Alain Garcia De Las Bayonas, Maxwell Coyle, Flora Rutaganira, Deepak Krishnamurthy, Samed Delic, Masayuki Onishi, Agathe Chaigne, and Ben Larson provided helpful feedback and suggestions for experiments. Samed Delic provided advice and helpful discussion on septin phylogenetics. Jacob Steenwyk provided advice and script construction for phylogenetic analysis. David Booth and Laura Wetzel provided valuable insights into previous work on choanoflagellate septins and relevant rosette-defective *S. rosetta* mutants. Stefany Gonzales aided with molecular cloning and choanoflagellate culture. Kevin Rose provided insights and suggestions for septin structural modeling. Maria Nguyen and Flora Rutaganira provided RIF-OMV cultures used throughout this work as well as guidance and equipment/reagent use for CRISPR/Cas9-mediated endogenous C-terminal fluorescent tagging in *S. rosetta*. M.D.C. was supported by an NIH Molecular Basis of Cell Function T32 Training Grant (5T32GM007232). An Investigator Award from the Howard Hughes Medical Institute supports research in the King Lab. This work used the UC Berkeley DNA Sequencing Facility (Molecular and Cell Biology Department).

## AUTHOR CONTRIBUTIONS

Michael D. Carver, Conceived and designed the experiments, Performed the experiments, Analyzed the data, Prepared the digital images, and Drafted the article | Nicole King, Conceived the experiments, Contributed methodology, Acquired funding, Supervised the work, and Drafted the article.

