## Supplemental Figures and Tables for "Septins regulate cytokinesis and multicellular development in the closest living relatives of animals"

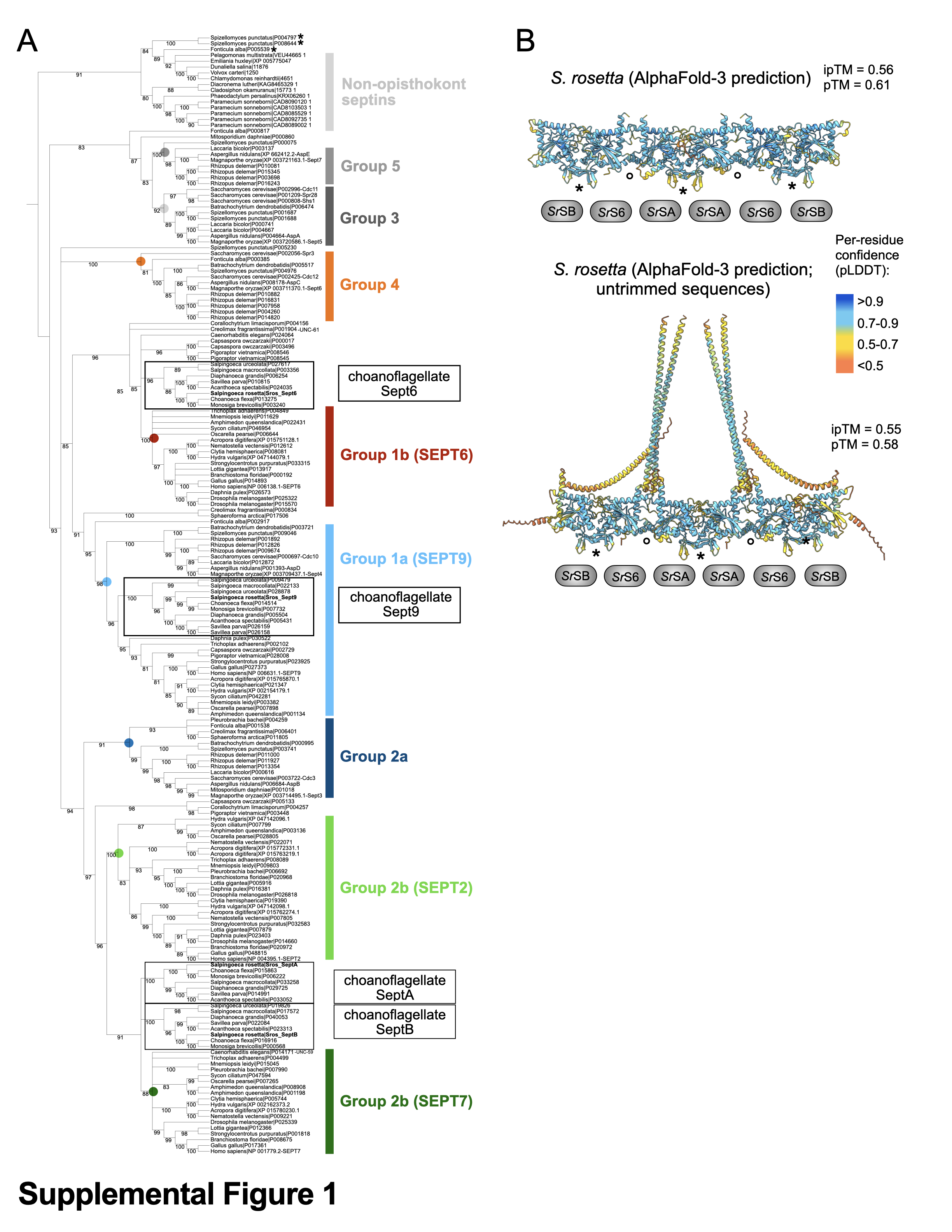

**Figure S1.** Phylogenetic relationships among diverse eukaryotic septins and predicted assembly of *S. rosetta* septins. (A) Choanoflagellate Sept9 confidently groups within the established fungal and animal SEPT9 clade (Group 1a), while choanoflagellate Sept6 forms a polytomy with animal SEPT6, septins from other close animal relatives, and the *C. elegans* septin *UNC-61* (Group 1b) (Shuman and Momany, 2022). Choanoflagellate SeptA and SeptB form a polytomy with animal SEPT7 within the broader SEPT2/7 animal clade (Group 2b). A maximum-likelihood phylogeny is shown of predicted septin protein sequences from diverse opisthokonts and selected non-opisthokont taxa (File S1). All nodes with less than 80% bootstrap support were collapsed prior to visualization. Established animal and fungal groups are indicated with colored bars and dots at the corresponding nodes, choanoflagellate septin groups are indicated with boxes, and *S. rosetta* septins in bold. The tree is rooted using non-opisthokont septins as an outgroup. Asterisks indicate septins from two early-branching holomycotans, which grouped within the non-opisthokont septins, potentially stemming from complex patterns of duplication and loss, horizontal acquisition, or analytical artifacts. (B) The *S. rosetta* septin hexamer is depicted with the predicted local distance difference test (pLDDT) scores, which largely fell in the 70 – 90 range (Varadi *et al.*, 2022; Abramson *et al.*, 2024). Global model confidence (pTM) and interface confidence (ipTM) values fall below commonly used confidence thresholds but are interpreted cautiously given known limitations with these metrics, including sensitivity to sequence trimming and known limitations in confidently predicting multimeric structures (Liang *et al.*, 2024; Dunbrack, 2025). Top, model generated from trimmed sequences to match the solved human crystal structure (Figure 1D); bottom, model generated without trimming and used to define alignment pairs between human and *S. rosetta* septins for sequence trimming. *S. rosetta* septins modeled: *Sros_*SeptB (*Sr*SB), *Sros_*Sept6 (*Sr*S6), and *Sros_*SeptA (*Sr*SA). Asterisks denote N- to C-terminal interfaces and circles denote GTP-binding domain interfaces. Residue-level confidence is shown as a color gradient based on predicted local distance difference test (pLDDT) scores. pTM and ipTM values are indicated for each structure.

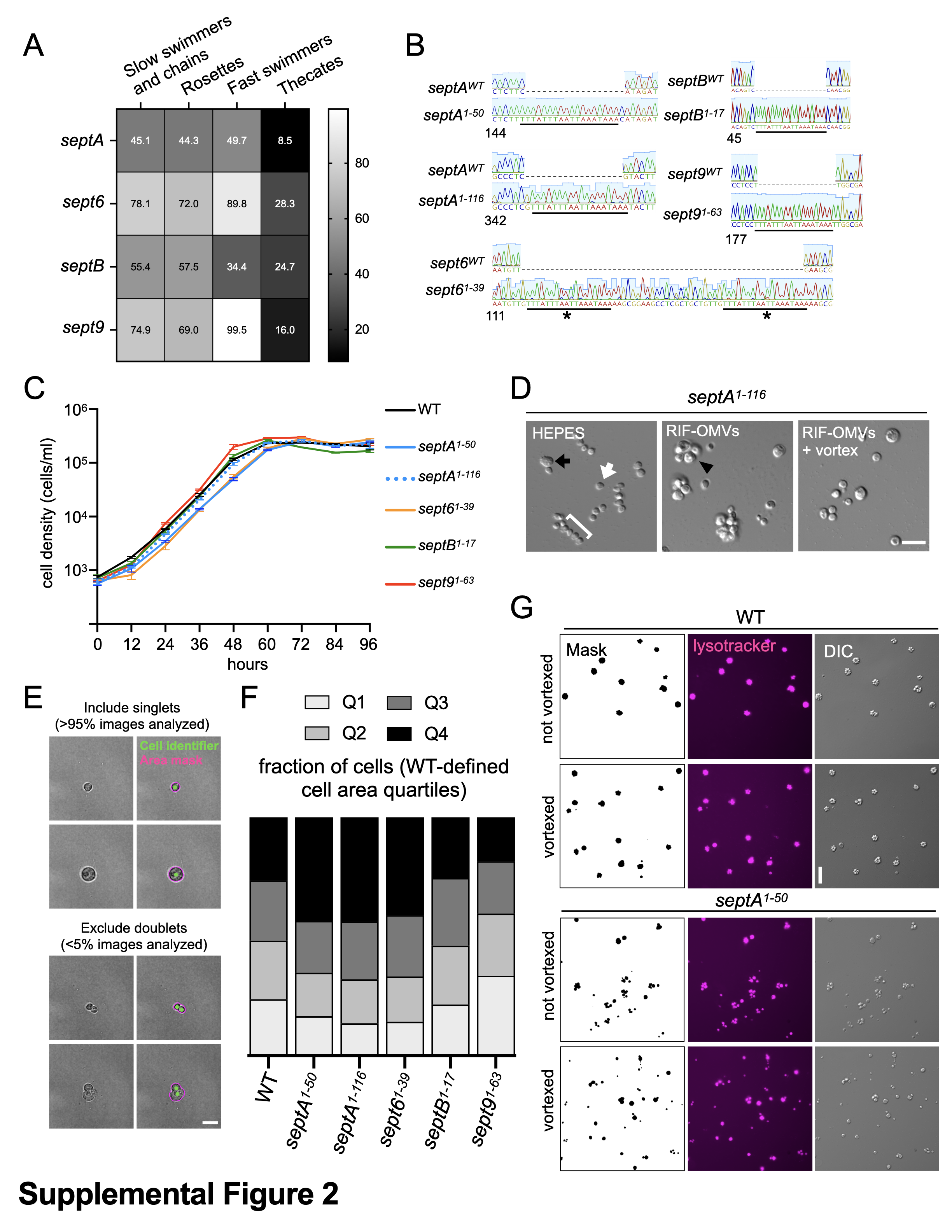

**Figure S2.** Structural modeling, expression, gene editing, and phenotyping of *S. rosetta* septins. (A) Relative to slow swimmers and chains, the expression of all four septins was significantly downregulated in thecate cells (q < 0.05 for all comparisons). In contrast, no significant differences were detected between slow swimmers and chains vs. rosette cells. Fast swimmers had a modest but significant reduction in *Sros*_*septB* expression (q = 0.048) compared with slow swimmers and chains, whereas the other three septins showed no significant differences. (B) A premature termination sequence was introduced into each of the four *S. rosetta* septin genes to create septin mutant alleles. Clonally isolated cells were edited with CRISPR/Cas9 and then genotyped by PCR and Sanger sequencing. The premature termination sequence TTTATTTAATTAAATAAA (black lines), encoding a stop codon in every possible reading frame, was introduced in an exon near the 5’ end of each gene. Numbers indicate position in the coding DNA sequence of the target gene. For *Sros*_*sept6^1-39^*, the premature termination sequence incorporated tandemly (asterisks). (C) Cell proliferation is slightly reduced in *Sros*_*septA^1-50^* and *Sros*_*sept6^1-39^* compared with wild-type. Cells were diluted to 5,000 cells/mL, triplicate samples were collected, and cell density was counted every 12 h for 96 h. Mean values were plotted and error bars denote the standard deviation. (D) Defects in cell size and rosette morphology in *Sros*_*septA^1-116^*. *Sros*_*septA^1-116^* cultures grown with a carrier control (HEPES) produced a mixture of single cells (white arrow) and chains (bracket) and also frequently contained larger single cells (black arrow). After treatment with RIF-OMVs, *Sros*_*septA^1-116^* cultures produced abnormal rosettes containing a mixture of normal and comparatively larger cells (black arrowhead). Following application of shear through vortexing, *Sros*_*septA^1-116^* cultures contained smaller colonies and more single cells compared with unvortexed *Sros*_*septA^1-116^* cultures, indicating a reduction in rosette integrity (quantification shown in Figure 2H). Scale bar, 20 μm. (E) Cell segmentation allowed for high-throughput analysis of projected area of single cells presented in Figure 2G. Cells analyzed through an imaging flow cytometer were processed through a cell segmentation pipeline to allow for accurate measurement of projected cell area. Single cells of variable sizes (top) were included in final analysis while images in which two or more cells were detected (bottom) were filtered out. The area mask (magenta) and cell identifier (green) are shown. Scale bar, 5 μm. (F) When grown without RIF-OMVs, *Sros*_*septA* and *Sros*_*sept6* mutants exhibited an increased frequency of oversized cells and a decreased frequency of smaller cells relative to wild-type. Conversely, *Sros_sept9^1-63^* exhibited an increased frequency of smaller cells and a decreased frequency of oversized cells. In all strains, the middle two quartiles remained similar. Cell area was quantified from individual cells pooled across three biological replicates (18,382 – 23,313 total cells per strain). Quartile ranges were defined using pooled wild-type data (Q1: <29.79 μm^2^; Q2: 29.79 – 35.73 μm^2^; Q3: 35.73 – 43.83 μm^2^; Q4: >43.83 μm^2^), and all septin mutant strains were analyzed relative to these thresholds. (G) An automated image analysis pipeline allowed for the quantification of rosette assembly and integrity phenotypes in wild-type and septin mutants shown in Figure 2H. Cells (top: differential interference contrast) were stained with LysoTracker Red DND-99 (middle) and imaged. By generating a binary mask from the fluorescent images, the projected area of each individual was measured (single cell, doublet, triplet, or rosette) in unvortexed and vortexed cultures, creating a proxy quantification of rosette assembly and integrity in response to shear, respectively. Representative images of wild-type and *Sros*_*septA^1-50^* are shown. Scale bar, 20 μm.

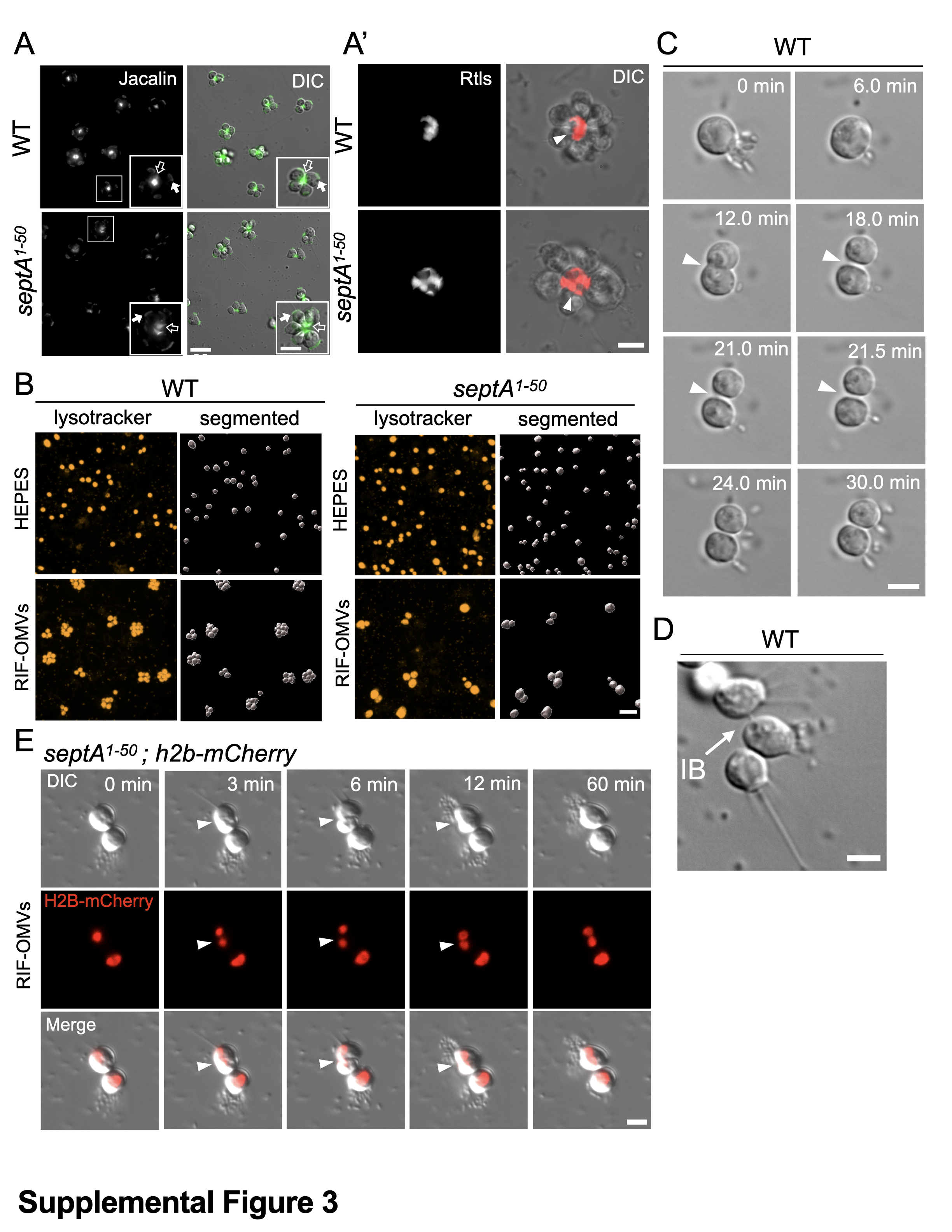

**Figure S3:** Cellular phenotypes of *Sros*_*septA* mutants and relevant wild-type *S. rosetta* cell biology. (A-A’) *Sros*_*septA^1-50^* cells secrete Jacalin-positive (A) and Rosetteless-positive (A’) ECM following rosette induction with RIF-OMVs. (A) Representative images of wild-type and *Sros*_*septA^1-50^* rosettes stained with fluorescein-labeled Jacalin (left), which stains the ECM of rosettes (empty arrow) and the collar base (filled arrow), along with a merged image that shows Jacalin-stained ECM along the basal cell poles at the center of each rosette. A representative rosette is highlighted in the inset for each strain. Scale bar, 10 μm. Inset scale bar, 5 μm. (A’) Representative maximum Z-projection images of wild-type and *Sros*_*septA^1-50^* rosettes stained with an antibody to Rosetteless protein (left), a component of the rosette ECM that is necessary for rosette formation (Levin *et al.*, 2014). Fluorescent images merged with DIC (right) show the ECM along the basal cell poles (arrowhead) at the center of each representative rosette. Scale bar, 5 μm. (B) A cell segmentation pipeline allowed for analysis of cell volume of wild-type (left) and *Sros*_*septA^1-50^* (right) single cells and rosettes. Three-dimensional confocal fluorescence imaging of single cells and rosettes stained with LysoTracker Red DND-99 (left, center Z-section is shown) followed by machine learning-based segmentation (right, 3D top-down renderings shown) allowed for accurate cell volume measurements of cells in rosettes, enabling comparison of cell size between uninduced and rosette-induced cultures in both wild-type (B) and *septA^1-50^* mutants (B’). Scale bar, 15 μm. (C) Wild-type cells undergo cytokinesis and remain attached following cell division. Times shown were chosen to match the visual cues (6 min – bacterial ejection from collar; 12 min – furrow ingression; 18 min – two cells appear distinct) of the *Sros_septA^1-50^* cytokinesis failure panels in Figure 3C. Furrow ingression at the basal pole is indicated (arrowhead). In this representative time series, unlike with *Sros_septA^1-50^*, a wild-type cell that divides between 6 and 18 min remains divided after 30 min (Video S1). Scale bar, 5 μm. (D) Cells in wild-type chains are connected by intercellular bridges. Shown is a representative image of a chain colony with an intercellular bridge (IB) indicated by a white arrow. Scale bar, 5 μm. (E) Cytokinesis failure in *Sros_septA^1-50^* results in multinucleated cells. A representative cytokinesis failure event is shown with a *Sros_septA^1-50^* cell expressing the nuclear marker H2B–mCherry following rosette induction: DIC (top); H2B–mCherry fluorescence (middle); merged images (bottom). H2B-mCherry-labeled nuclei divide during mitosis and remain distinct following cytokinesis failure, resulting in a multinucleated cell (Video S3). White arrowheads indicate the cleavage furrow. Time (minutes) is indicated. Scale bar, 5 μm.

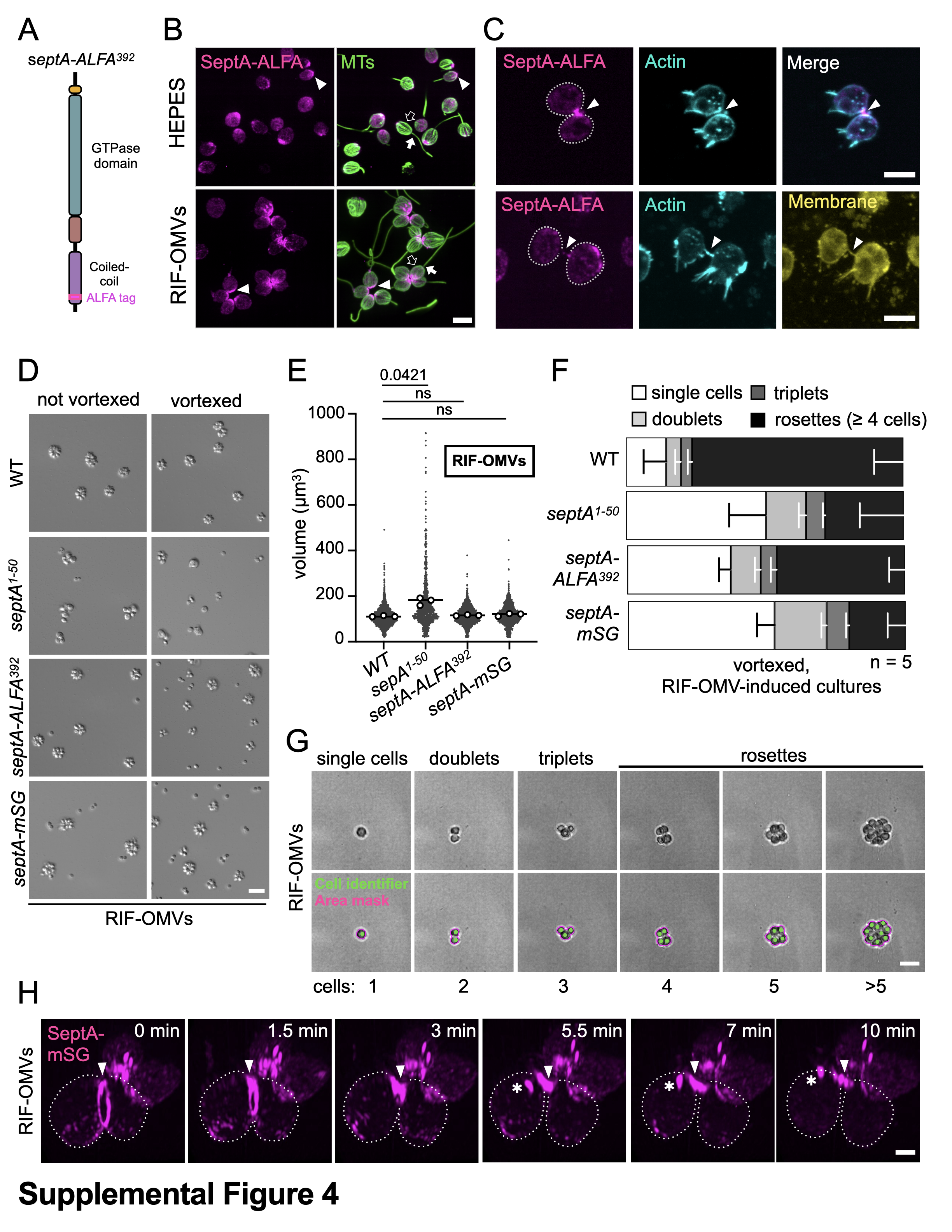

**Figure S4.** Localization and phenotypic effects in ALFA-tagged and mStayGold-tagged *Sros_septA* strains. (A) *Sros*_*septA* was endogenously tagged with an ALFA epitope at amino acid position 392 of the predicted protein sequence, enabling immunofluorescence imaging using anti-ALFA nanobodies (Götzke *et al.*, 2019). (B) SeptA-ALFA^392^ localization patterns are similar to those of SeptA-mStayGold (Figure 4B). *Sros*_SeptA-ALFA stained with an anti-ALFA nanobody (left) reveals cytoplasmic and basal enrichment (arrowhead), while microtubules marking the cell body (“tubulin cage”, empty arrow) and apical flagella (filled arrow) were visualized with an anti-⍺-tubulin antibody. Scale bar, 5 μm. (C) *Sros_*SeptA-ALFA localizes to intercellular bridges (arrowhead) in dividing cells. Top: representative example of strong SeptA-ALFA localization. Bottom: representative example of weak SeptA-ALFA localization. Actin is labeled with phalloidin; membrane is labeled with FM-1-43-FX dye. Scale bar, 5 μm. (D) Rosette integrity defects are associated with tagging *Sros_septA*. Representative images of rosette-induced cultures before and after application of shear through vortexing are shown. Wild-type rosettes appear unaffected after shear, while *Sros*_*septA^1-50^* rosettes disassemble. *Sros*_*septA-ALFA^392^* and *Sros_septA-mSG* strains also disassemble, with more single cells and smaller rosette structures following the application of shear. Scale bar, 10 μm. (E) Tagged septin strains do not have a cell size defect following rosette induction. In contrast to *Sros*_*septA^1-50^*, mean cell volume is not significantly different in rosette-induced tagged septin strains compared with wild-type. Dots represent cell volume of individual cells pooled from three biological replicates following 24 h RIF-OMV treatment (831–1528 total cells); white dots represent means per replicate; bars denote means across replicates. Statistics performed on replicate means (n = 3): one-way ANOVA with Dunnett’s multiple comparisons test to compare between strains**.** (F) Rosette integrity is decreased in tagged septin strains compared with wild-type. Using the cell segmentation pipeline described in (G), the fraction of individuals that were single cells, doublets, triplets, or rosettes per strain were determined and the means of 5 biological replicates (25,000 cells pooled in total) per strain are plotted with error bars representing standard deviation. (G) Image analysis pipeline allowed for high-throughput analysis of cells per individual following vortexing. Vortexed, rosette-induced cultures passed through a flow cytometer were imaged and processed through a cell segmentation pipeline that reliably identified cells per individual for structures up to 5 cells. Individual mask is shown in magenta while the cell identifier is in green. Scale bar, 10 μm. (H) SeptA-mSG accumulates in a ring at the cleavage furrow (filled arrowhead) and nascent intercellular bridge of dividing rosette-induced cells (Video S5) (localization in uninduced dividing cells shown in Figure 4C). As cleavage progresses, the SeptA-mSG ring constricts and becomes enriched at the nascent intercellular bridge connecting the daughter cells (open arrowhead), resolving into two puncta after several minutes. Dotted lines outline cell boundaries. Scale bar, 1 μm. In both single cells and rosette cells, we additionally detected dynamic SeptA-mSG puncta of unknown function (Figure 4C and S4H, asterisks).

**SUPPLEMENTAL TABLES**

|  | Table S1 — Comparison of *S. rosetta* septin expression across life history states | | | | | |
| --- | --- | --- | --- | --- | --- | --- |
| **Septin** | | **Life history stage** | **Fold**  **change vs slow swimmers/chains** | **p-value** | **q-value** | **Interpretation** |
| *Sros*_*septA* | | rosette | **-0.036** | **0.859** | 1.000 | Not significantly different |
| *Sros*_*sept6* | | rosette | **-0.121** | **0.539** | 1.000 | Not significantly different |
| *Sros*_*septB* | | rosette | **0.063** | **0.802** | 1.000 | Not significantly different |
| *Sros*_*sept9* | | rosette | **-0.126** | **0.527** | 1.000 | Not significantly different |
| *Sros*_*septA* | | fast swimmer | **0.139** | **0.496** | 0.669 | Not significantly different |
| *Sros*_*sept6* | | fast swimmer | **0.204** | **0.302** | 0.498 | Not significantly different |
| *Sros*_*septB* | | fast swimmer | **-0.657** | **0.008** | 0.048 | Significant, slight decrease |
| *Sros*_*sept9* | | fast swimmer | **0.415** | **0.036** | 0.129 | Not significantly different |
| *Sros*_*septA* | | thecate | **-2.453** | **1.95e-33** | 2.58e-32 | Significantly lower |
| *Sros*_*sept6* | | thecate | **-1.506** | **2.59e-14** | 1.15e-13 | Significantly lower |
| *Sros*_*septB* | | thecate | **-1.208** | **1.26e-6** | 3.04e-6 | Significantly lower |
| *Sros*_*sept9* | | thecate | **-2.273** | **1.00e-30** | 1.17e-29 | Significantly lower |

|  | Table S2 — Mean cell size (slow swimmers & chains) | | | | |
| --- | --- | --- | --- | --- | --- |
| **Strain** | | **Mean cell area (µm²)** | **Fold**  **change** | **p-value vs WT** | **Interpretation** |
| **Wild-type** | | **41.28** | — | — | Reference |
| *Sros*_*septA^1-50^* | | 51.75 | **1.25**× | **< 0.0001** | Significantly larger |
| *Sros*_*septA^1-116^* | | 49.28 | **1.19**× | **0.0002** | Significantly larger |
| *Sros*_*sept6^1-39^* | | 46.77 | **1.13**× | **0.005** | Significantly larger |
| *Sros*_*sept9^1-63^* | | 37.15 | **0.90**× | **0.0286** | Significantly smaller |
| *Sros*_*septB^1-17^* | | 38.91 | 0.94× | 0.2684 | Not significantly different |

**Table S3 — Mean projected “individual” size after RIF-OMV–induced rosette formation** (Individuals = single cells, doublets, triplets, or rosettes)

| **Strain** | | **Mean area (µm²)** | **p-value vs WT** | **Interpretation** |
| --- | --- | --- | --- | --- |
| **Wild-type** | | **168.3** | — | Reference |
| *Sros*_*septA^1-50^* | | 110.1 | **0.0002** | Significantly smaller |
| *Sros*_*septA^1-116^* | | 102.1 | **< 0.0001** | Significantly smaller |
| *Sros*_*sept6^1-39^* | | 90.2 | **< 0.0001** | Significantly smaller |
| *Sros*_*sept9^1-63^* | | 107.9 | **< 0.0001** | Significantly smaller |
| *Sros*_*septB^1-17^* | | 157.8 | 0.8236 | Not significantly different |

**Table S4 — Mean projected individual size after RIF-OMV-induced formation *after exposure to shear*** (Individuals = single cells, doublets, triplets, or rosettes)

| **Strain** | **Mean area after shear (µm²)** | **p-value (vs pre-shear)** | **Interpretation** |
| --- | --- | --- | --- |
| **Wild-type** | **157.9** | 0.9382 | Rosettes intact |
| *Sros*_*septA*^1-50^ | 77.05 | **0.0465** | Significant disassembly |
| *Sros*_*septA*^1-116^ | 68.48 | **0.0421** | Significant disassembly |
| *Sros*_*sept6*^1-39^ | 56.03 | **0.0373** | Significant disassembly |
| *Sros*_*sept9^1-63^* | 100.2 | 0.9856 | No significant disassembly |
| *Sros*_*septB^1-17^* | 141.0 | 0.6301 | No significant disassembly |

**Table S5 — Effect of RIF-OMV-induced rosette induction on cell volume**

| **Strain** | **Condition** | **Mean cell volume (µm³)** | **Fold change** | **p-value** | **Interpretation** |
| --- | --- | --- | --- | --- | --- |
| **Wild-type** | Uninduced | 140.2 | — | — | Reference |
|  | RIF-OMV | 115.7 | 0.83× | 0.8345 | No significant change |
| ***Sros*_*septA^1-50^*** | Uninduced | 145.9 | — | — | Reference |
|  | RIF-OMV | 515.1 | **3.53×** | **< 0.0001** | Strong increase |
| ***Sros*_*septA^1-116^*** | Uninduced | 162.2 | — | — | Reference |
|  | RIF-OMV | 250.0 | **1.54×** | **0.0485** | Moderate increase |

**Table S6 — Cytokinesis failure rates during the first 10 hours following RIF-OMV rosette induction**

| **Strain** | **Condition** | **Cytokinesis failure rate (%)** | **Comparison** | **p-value** | **Interpretation** |
| --- | --- | --- | --- | --- | --- |
| **Wild-type** | Uninduced | 2.0 | — | — | Baseline |
| **Wild-type** | RIF-OMV | 4.6 | vs WT uninduced | 0.1101 | No significant induction effect |
| ***Sros*_*septA^1-50^*** | Uninduced | 12.2 | vs WT uninduced | 0.0292 | Elevated vs WT |
| ***Sros*_*septA^1-50^*** | RIF-OMV | 30.9 | **vs *Sros_septA^1-50^* uninduced** | **0.0115** | Significantly higher failure rate following induction within ***Sros*_*septA^1-50^*** |

**SUPPLEMENTAL VIDEOS**

**Video S1. Wild-type S. rosetta cells typically complete cytokinesis.**

DIC time-lapse microscopy of a wild-type S. rosetta cell undergoing cell division. Time (hours:minutes:seconds) is indicated. Scale bar, 2 µm. Related to Figure S3C.

**Video S2. *Sros*_*septA^1-50^* mutant cells frequently fail cytokinesis following RIF-OMV induction.**

DIC time-lapse microscopy of a *Sros_septA^1-50^* cell following rosette induction with RIF-OMVs. After cleavage furrow ingression, the two daughter cells fuse, indicating a defect in a late stage of cytokinesis. Time (hours:minutes:seconds) is indicated. Scale bar, 2 µm. Related to Figure 3C.

**Video S3. Nuclei remain separated after cells re-fuse in *Sros*_*septA^1-50^* mutants.**

DIC and fluorescence time-lapse microscopy of *Sros*_*septA^1-50^* cells expressing the nuclear marker H2B–mCherry following rosette induction with RIF-OMVs. Left: merge of H2B–mCherry fluorescence and DIC. Right: H2B–mCherry fluorescence. H2B–mCherry-labeled nuclei divide during mitosis and remain distinct following cytokinesis failure, resulting in a large, multinucleated cell. Time (hours:minutes:seconds) is indicated. Scale bar, 5 µm. Related to Figure S3E.

**Video S4. SeptA-mStayGold localizes to the cleavage furrow in uninduced dividing cells.**

Three-dimensional fluorescence time-lapse microscopy of an endogenously tagged SeptA-mStayGold (Sros_septA-mSG) cell dividing under uninduced conditions (slow swimmer). SeptA-mSG accumulates in a ring at the cleavage furrow during ingression, constricts as cytokinesis progresses, and becomes enriched at the nascent intercellular bridge, ultimately resolving into two puncta. Time (hours:minutes:seconds) is indicated. Related to Figure 4C.

**Video S5. SeptA-mStayGold localizes to the cleavage furrow in rosette-induced dividing cells.**

Three-dimensional fluorescence time-lapse microscopy of an endogenously tagged SeptA-mStayGold (Sros_septA-mSG) cell dividing following rosette induction with RIF-OMVs. Similar to uninduced cells, SeptA-mSG accumulates at the cleavage furrow and nascent intercellular bridge, resolving into two puncta after several minutes. Scale bar, 1 µm. Related to Figure S4H.

**SUPPLEMENTAL DATA**

**File S1. Data underlying phylogenetic and structural analysis of septins.**

This file documents the identification and filtering of septin sequences used to construct the phylogeny in Figure S1A. Tab “probe_sequences" lists the four human septin sequences (SEPT2, SEPT6, SEPT7, SEPT9) used as BLAST probes. Tab "species_used" lists all species queried. Tab "interproscan_putseptins_all" provides raw InterProScan domain predictions for all 201 candidate sequences. Tab "PF00735-positives" lists the protein length and lowest e-value PF00735 domain for each of the 201 putative septins. Tab "filtered_final_list" contains the 192 sequences retained after filtering by domain architecture and protein length. Tab "excluded" lists the 9 sequences removed after domain filtering, with reasons. Tab “structural analysis” contains the N- and C-terminally trimmed human septins used to align and trim the *S. rosetta* septins (also listed) used for AlphaFold-3 structural predictions.

**File S2. Corrected Sros_septB (PTSG_04364) coding and protein sequence.**

Tab "Sros_septB_corrected-seq" provides the revised Sros_septB CDS and protein sequences, corrected using RNA-seq read evidence (deposited at GenBank under accession PZ120989). Tab "RNAseq_reads_revision" documents the RNA-seq read support counts used to identify and correct intron–exon boundaries in the original genome annotation, with intron coordinates, strand, and read support for each junction.

**File S3. Differential expression of S. rosetta septin genes across cell states.**

Tab "Sros_septins_RNAseq" reports pairwise differential expression statistics for all four S. rosetta septin genes (Sros_septA, Sros_sept6, Sros_septB, Sros_sept9) across cell state comparisons (slow swimmers and chains vs. thecate, rosette, and fast swimmer cells). The upper section includes TPM values for individual biological replicates (three per condition), condition averages, standard deviations, p-values, q-values, log₂ fold-change, and standard error of fold-change, as well as Ensembl, UniProt, Pfam, InterPro, GO, and eggNOG annotations. The lower section summarizes results using condition averages only, with a qualitative "Comment" column. Data are from Leon *et al*. (2025) and underlie Figure S2A and Table S1.

**File S4. CRISPR-Cas9 reagents for septin gene disruption and tagging.**

Tab "CRISPR gene edits" provides gRNA sequences, repair template sequences, insertion sites (CDS position), and genotyping primer pairs for all gene truncation and tagging experiments, including alleles of Sros_septA (two truncation lengths: 50 and 116 aa; plus ALFA-tag insertion), Sros_sept6 (39 aa), Sros_septB (17 aa), Sros_sept9 (63 aa), and the rpl36a-P56Q cycloheximide-resistance locus. Also includes editing efficiency metrics (number of experiments, clones genotyped, and independent edited clones obtained). Tab "ALFA-tag" provides further details about the ALFA-tag repair template design, including annotated sequences for the crRNA binding site sequence and ALFA-tag insertion, along with the associated *Sros*_*septA* CDS.

**File S5. Reagents for endogenous Sros_SeptA–mStayGold (SeptA-mSG) tagging.**

Tab "SeptA-mSG_sequences" documents all sequences used to generate the SeptA-mSG knock-in strain (NK812), including: the mStayGold nucleotide and protein sequences; the SGGSGGS linker sequences; the gene block for Gibson assembly into plasmid pMS18; Gibson cloning and PCR amplification primers (with annealing temperature and extension time); the crRNA sequence; and three genotyping primer pairs for confirming correct insertion. Tab "SeptA-mSG_codon-changes" lists the eight synonymous codon substitutions introduced into the plasmid-encoded Sros_septA recovery region to reduce homologous recombination with the repair template, with original and replacement codons and their relative usage frequencies (from Leon *et al*. (2025)).
